# Hormone-regulated dynamics of mRNA distribution on ribosomes in Sertoli cells

**DOI:** 10.1101/2024.12.20.629416

**Authors:** Xavier Leray, Julia Morales, Aurélie Tréfier, Kelly León, Thomas Bourquard, Astrid Musnier, Emmanuel Pecnard, Lucile Drobecq, Sandrine Boulben, Nicolas Azzopardi, Rémi Coulaud, Hamza Ouchène, Yan Jaszczyszyn, Eric Reiter, Florian Guillou, Lucie P. Pellissier, Anne Poupon, Romain Yvinec, Pascale Crépieux

**Author notes:** To whom correspondence should be addressed: Dr. Pascale Crépieux, UMR Physiologie de la Reproduction et des Comportements, Centre de Recherches INRAE Val de Loire, F-37380 Nouzilly, France. Dr. Romain Yvinec, UMR Physiologie de la Reproduction et des Comportements, Centre de Recherches INRAE Val de Loire, F-37380 Nouzilly, France. Co-senior authors (Romain Yvinec, Pascale Crépieux).

## Abstract

The effects of hormone stimulation on the cell translational profile remain poorly understood. Here, using polysome profiling combined to RNA sequencing, we analyzed the translational response to follicle-stimulating hormone (FSH) of primary rat Sertoli cells, that exhibit an active anabolic activity regulated by reproductive hormones in the male gonad. We first established that mRNA distribution to polysomes follows a bimodal pattern, with 15% of mRNAs enriched in polysomes and exhibiting high expression. Critically, this basal polysomal enrichment had a major impact on FSH-induced mRNA recruitment to the polysomes, since FSH stimulation promoted the release of polysome-enriched mRNAs, while mRNAs that were the least associated to polysomes were preferentially recruited to polysomes upon stimulation. The FSH signal did not alter the core biological functions of Sertoli cells, but shifted the proteins involved in these functions, suggesting a molecular rewiring of the FSH-induced gene expression. These findings underscore how ribosomal reallocation dynamically adapts the cellular translatome to microenvironmental changes, enabling cells to fine-tune protein production in response to external stimuli.

**GRAPHICAL ABSTRACT:** 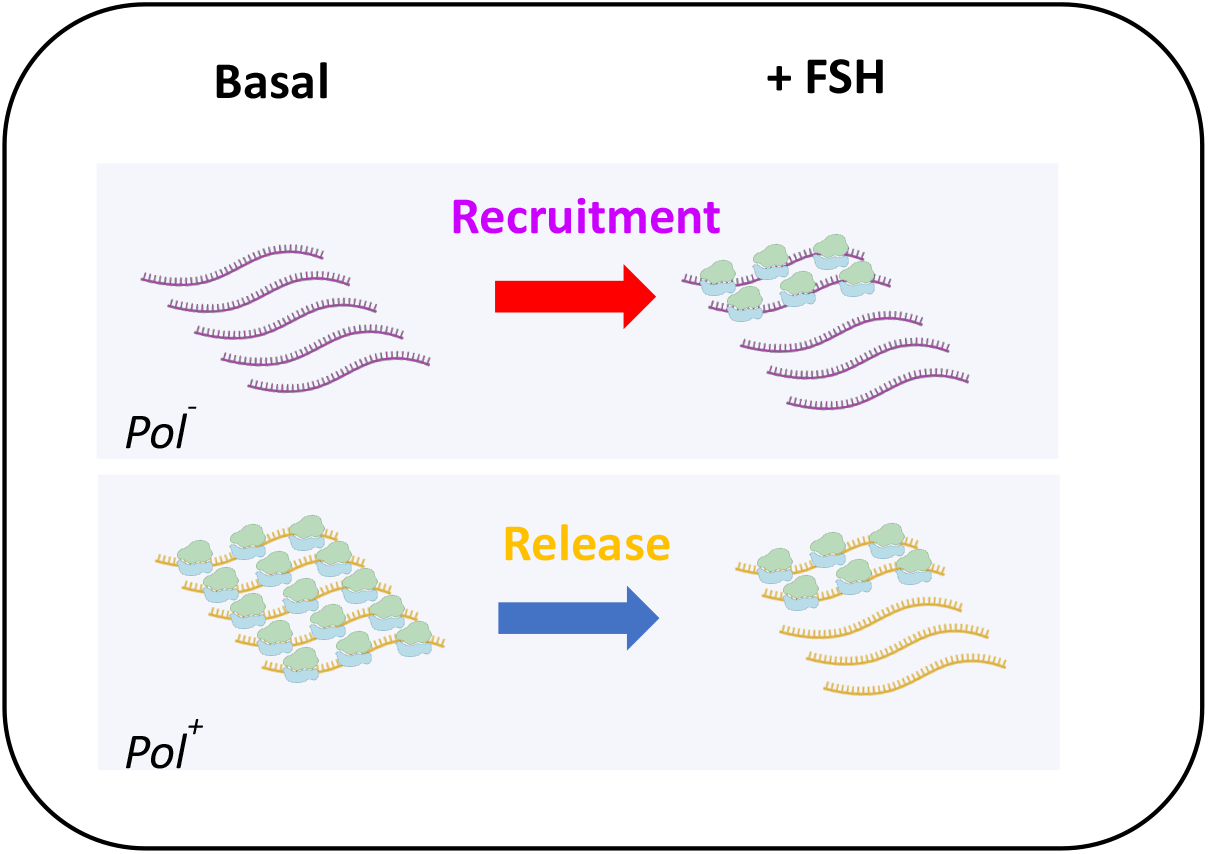

**Bullet points:** • In Sertoli cells, most mRNAs distribute similarly between monosomes and polysomes, but a sub-population is specifically enriched in polysomes

• Basal polysomal enrichment level has a major impact on FSH-induced mRNA recruitment or release from the polysomes

• The FSH signal induced a global rewiring of the proteins involved in Sertoli cell basal activity

• FSH-induced reassignment of ribosomes to specific mRNAs has to comply with a tightly maintained mRNA distribution landscape

## Introduction

To fulfill its physiological roles, a cell continuously senses and responds to changes in its surrounding environment by adjusting the type, quantity and activity of its expressed proteins. This precise regulation of protein copy number is achieved through the balance between synthesis and decay reactions (Hargrove and Schmidt, 1989; Li and Biggin, 2015; Liu et al., 2016). Protein synthesis results from transcription and translation, the latter being influenced by the number of ribosomes allocated per molecule of mRNA (Riba et al., 2019). Simply put, a higher ribosomal load increases protein synthesis, hence, adjusting the ribosomal loading is a key mechanism to adapt protein synthesis to environmental changes (Liu et al., 2016; Sonenberg and Hinnebusch, 2009).

Transcriptional (Jovanovic et al., 2015; Li et al., 2014; Li and Biggin, 2015) and translational control (Schwanhäusser et al., 2011) have been debated as primary driver of the cellular protein content, although these processes are essentially uncoupled (Tebaldi et al., 2012). While transcription reflects steady-state protein levels, translation plays a more dynamic role, enabling rapid adaptation to extra-cellular signals or environmental changes (Liu et al., 2016).

In polysome profiling studies, cytosolic mRNAs classically distribute into 3 pools with distinct ribosomal loading (Chassé et al., 2017): (1) the free pool (Free) where mRNAs are not bound to ribosomes, (2) the monosomal pool (Mono) where mRNAs bind a single ribosome, (3) and the polysomal pool (Pol) where mRNAs are associated to at least two ribosomes and are actively translated. While recent studies have explored the role of Mono in mRNA local translation (Biever et al., 2020; Heyer and Moore, 2016) and while the role of Free remains enigmatic (Arribere et al., 2011; Luo et al., 2018; Wang et al., 2018), most translatomic studies focus on changes within Pol. In actively translating cells, most mRNAs are ribosome-bound, with ribosome occupancies -defined as the fraction of transcripts bound by at least one ribosome- varying between 50% to >90% across various species, including yeast, plant and bacterium (Arava et al., 2003; Kawaguchi and Bailey-Serres, 2005; Lackner et al., 2007; Nguyen et al., 2022; Picard et al., 2012). However, whether the basal distribution of mRNAs among Free, Mono and Pol influences their recruitment to polysomes during cell stimulation remains unclear. For example, it is unknown if mRNAs highly enriched in polysomes can be further recruited upon cell stimulation. If the basal distribution does impose constraints, changes in mRNA levels within polysomes may reflect pre-existing distribution patterns rather than purely the effects of signaling mechanisms. This consideration is of great interest for translatomic studies, since such a constraint may lead to misinterpret polysomal changes as merely originating from the signaling mechanisms induced by the external stimulus.

Adapting to environmental variations involves intricate mechanisms of cellular plasticity. For instance, IFN𝛾-mediated mTOR inhibition induces profound translational changes that modulate inflammatory responses and autophagy in immune cells (Su et al., 2015). Yet, translatomic studies addressing the effects of extra-cellular signals remain limited. In particular, hormones represent key extracellular signals. Many of them bind G protein-coupled receptors (GPCR), and the translatome of very few of the 800 known GPCRs has been reported, whether investigated by polysome profiling or by in vivo methods like Translating Ribosome Affinity Purification (TRAP) (Tréfier et al., 2018c, 2018a). Notable examples include the translatomes of mGluR1/5 (Di Prisco et al., 2014), GnRHR (Do et al., 2014) and ET1R (Markou et al., 2010). Recent mechanistic details on GPCR-mediated translation have come from ribosome profiling of the β2-adrenoceptor activity, which demonstrates endosome-localized receptor-mediated translational changes, predominantly affecting 5’TOP mRNAs (Klauer et al., 2024).

In the reproductive tract, analysis of Leydig cell-specific RiboTag mice have identified a small subset of luteinizing hormone (LH)-responsive mRNAs at the translational level, though distinction between monosomal and polysomal-bound mRNAs was not addressed (Sanz et al., 2013). In the testis, follicle-stimulating hormone (FSH) supports spermatogenesis by stimulating Sertoli cell mitoses in neonate and by sustaining their secretory activity in adulthood (Meachem, 1996; Orth, 1984; Santi et al., 2020; Zimmermann et al., 2015). FSH acts through the FSH receptor (FSHR), a GPCR located at the basal membrane of Sertoli cells. FSHR activation initiates a signaling network that ultimately affects the copy number and activity of specific proteins, driving Sertoli cell anabolic activity to nurture germ cells and cytoskeleton remodeling essential for the blood-testis barrier dynamics (Ulloa-Aguirre et al., 2018). While several transcriptomic studies have provided insights into FSH-induced long-term transcriptional changes (Abel et al., 2009; Crespo et al., 2016; McLean et al., 2002; Meachem et al., 2005; O’Donnell et al., 2009; Sadate-Ngatchou et al., 2004), less is known about short-term translational regulations. Evidence have shown that the FSHR signaling network targets the translational machinery via mTOR signaling (León et al., 2014; Musnier et al., 2009, 2012), transduced by G proteins and β-arrestins (Tréfier et al., 2018b). However, the specific identities of regulated mRNAs and the biological processes they govern remain largely unexplored. In this study, our goal was to uncover the general principles underlying mRNA distribution to polysomes, and to gain insights into the translational regulation orchestrated by FSH.

## Materials and Methods

References of the materials are provided in **Supplementary Table S1**.

### Experimental procedure

#### Isolation and culture of primary Sertoli cells

Sertoli cells were isolated from testes of 19-day-old Wistar rats from different littermates (Iffa- Credo, Lyon, France). Animals were treated following the current ethical guidelines of the European Community 86/609/CEE. Following purification as reported previously (Guillou et al., 1986), Sertoli cells (15 x 10^6^ cells per condition) were seeded on CellBind plates (Corning Life Sciences) in DMEM (Eurobio Scientific) complemented with penicillin (100 IU/ ml) and streptomycin (100 µg/ml) manufactured by Gibco, and with glutamine (2 mM) from Eurobio Scientific. Retinol (50 ng/ml), vitamin E (200 ng/ml) and human transferrin (5 µg/ml), all purchased from Sigma, were also added. In average, our Sertoli cell cultures were 90 % pure, as previously quantified (Guillou et al., 1986). Cells were maintained in a humidified atmosphere with 5% CO_2_ at 34°C.

#### Polysomal fractionation on sucrose density gradient and RNA purification

Polysome profiling was achieved as reported in (Musnier et al., 2012). Forty-eight hours after initial seeding, the medium was changed and cells at 80 % confluence were exposed for 90 minutes to 100 ng/ ml porcine FSH, obtained from Dr George R. Bousfield (Wichita State University, KS, USA), or were left untreated. This short time of cell stimulation was chosen to minimize transcriptional responses, which are estimated to peak at 6 hrs in response to FSH (McLean et al., 2002). Fifteen minutes before harvesting, 100 μg/ml of cycloheximide (CHX, from Sigma) was added. Then, cells were washed with ice cold PBS (Dulbecco)/CHX and scraped in polysome lysis buffer [10 mM Tris-HCl (Pufferan) pH 7.8, 10 mM MgCl_2_ (Carlo Erba), 1.5 mM KCl (Sigma), 1 % Igepal-CA630 (Sigma), 1 % deoxycholate (Carol Roth GmbH), 2.5 mM dithiothreitol (Pierce), 100 μg/ml CHX] and 500 U/ml RNasin (Promega). Cellular debris were removed by centrifugation at 10, 000 rpm for 5 min and supernatants were layered onto a 12 ml linear sucrose gradient (10-45 % sucrose in 10 mM Tris-HCl pH 7.8, 10 mM MgCl_2_, 80 mM KCl and 1 mM DTT) and centrifuged at 38, 000 rpm in an SW41-Ti rotor at 4°C for 150 min. Using a density gradient fractionation system (ISCO, Inc.), 21 fractions of 600 𝜇l each were collected while absorbance was monitored at 254 nm. Fractions were treated with 200 µg/ml proteinase K (Eurobio Scientific) for 25 min at 50°C in 10 mM Tris-HCl pH 8.0, 1 mM EDTA (Sigma), 0.5 % SDS (Sigma), and then deproteinized with an equal volume of phenol: chloroform (1: 1) and precipitated with isopropanol. The RNA from each fraction was dissolved in 30 μl diethyl pyrocarbonate (Sigma)-treated H_2_O, and an aliquot was subjected to electrophoresis through a 1 % agarose gel, stained with GelRed (Interchim). A control experiment was done as above, except that the samples were pretreated with 50 mM EDTA for 10 min prior to loading and that the linear sucrose gradient was supplemented with 10 mM EDTA.

#### Construction of cDNA libraries and RNA sequencing

The following steps were performed at the Institute for Integrative Biology of the Cell (I2BC) (Gif-sur-Yvette, France). The RIN (RNA Integrity Number) was estimated for each sample using an Agilent Bioanalyzer Pico chip. cDNA libraries were constructed by using the TotalScript RNA-Seq kit from Epicentre (discontinued), a transposon-based method of tagmentation (simultaneous tagging and fragmentation) that permits amplification from less than 5 ng of total RNA. For synthesis of the first strand, oligodT priming was chosen to limit rRNA amplification. The library products (inserts of 200 bp in average) were sequenced using an Illumina HiSeq 1000 equipment, as single-end 1 x 50 nucleotide reads. The sequencing yield post-trim ranged between 16 and 85 million reads per sample.

### Data analysis

#### Data availability

All the mRNA lists that support the findings of this study are available as Supplementary Tables referenced in the main text. The raw data are deposited at ZENODO and access will be free upon publication.

#### RNA-seq quality control

Analysis was performed on the sequencing data with FastQC (version 0.10.1) (http://www.bioinformatics.bbsrc.ac.uk/projects/fastqc/). Sequences in .bam format were aligned to Rattus norvegicus genome v6.08 using the Python package HTSeq (http://www-huber.embl.de/users/anders/HTSeq/doc/overview.html). Alignment parameters were chosen as follows: stranded = no; minaqual (% mismatch) = 10; feature type = transcript; intersection non-empty. Multiple alignments were discarded, as well as ambiguous assignations.

Replicate quality was assessed by inspection of raw counts distributions, clustering and principal component analysis. The analysis revealed that one replicate in the non-stimulated Pol condition and one replicate in the FSH-stimulated Free condition (samples 7 and 10) were largely different from any other replicates (**Supplementary Table S2**), and were discarded of the final analyses.

#### RNA-seq filtering

After removal of non-mRNA biotypes, the subsequent analyses were performed on a filtered gene list as follows: mRNAs without known gene symbol were removed; when duplicated, the mRNAs corresponding to a single gene symbol were summed; mRNAs that had at least 1 CPM in at least 2 replicates, corresponding to 10-15 reads per mRNA in the total pool and 4-7 in the translatome, were filtered in. This first round of filtering recovered 17,929 genes. Finally, mRNAs that were not recognized by the StringApp (see below the ‘*Gene set functional analysis’* section) were filtered out, leading to a final list of 14,646 genes.

#### Statistical analysis

Most statistical analyses were performed using R (https://www.r-project.org). Essentially, two methods, namely edgeR and DEseq2, were used to compare the RNA pools (free, monosomes, polysomes) in FSH-stimulated and non-stimulated conditions, and provided similar results (data not shown). Based on the 14,646 mRNAs of the STRING list within the remaining 16 samples (**Supplementary Table S2)**, principal component analysis was performed using DESeq2 with the variance stabilized counts data in a blind manner, that did not used the data design information.

#### mRNA distribution between Free, Mono and Pol

mRNA distributions among the free, monosomes and polysomes were assessed using standard DESeq2 (Love et al., 2014) differential analysis. The corresponding mRNA pools were designed as Free, Mono and Pol throughout the text. Specifically, unstimulated Free, Mono and Pol replicates were normalized together using DESeq2 function with the fraction type as data design. Fold-change without shrinkage correction and with adjusted p-value were subsequently deduced with the same function, using two by two comparisons (Free *versus* Mono, Free *versus* Pol, Mono *versus* Pol). The gene lists were deduced from these statistical tests as follows: Pol^+^ are mRNAs which are statistically more abundant in Pol when compared with each of the two other fractions (Free *versus* Pol and Mono *versus* Pol; adjusted p-value <0.05). The same applies for Mono^+^ and Free^+^. Pol^-^ are mRNAs which are statistically less abundant in Pol when compared to each of the two other fractions. The same applies for Mono^-^ and Free^-^. A similar methodology was applied to the FSH-stimulated Free, Mono and Pol samples. Calculation of Pol, Mono and Free proportions were obtained using normalized Pol, Mono and Free values divided by the sum of these three normalized pool values (Pol + Mono + Free).

#### Normalization of the effect of FSH

The effect of the FSH hormonal stimulation was assessed on each fraction independently. Specifically, for each fraction, unstimulated and stimulated replicates were normalized together and differential expression was assessed using the DESeq2 function, as described above. In particular, the Pol_REG_ mRNA list was defined as the mRNAs statistically less or more abundant (p-value <0.05) in Pol after FSH stimulation. The same applies for Mono_REG_ and Free_REG_.

#### Gene set functional analysis

The 14,646 identified mRNA set and its different subpopulations were analyzed in Cytoscape (v3.10.1) (Shannon et al., 2003) with the following app versions: StringApp (v2.0.1) (Doncheva et al., 2019), EnrichmentMap (v3.3.6) (Merico et al., 2010), clusterMaker2 (v2.3.4) (Morris et al., 2011), yFiles layout Algorithms (v1.1.3).

#### Tissue gene set enrichment analysis

The 14,646 identified mRNA set obtained from Sertoli cells was compared to the rat genome using the *functional enrichment tool* of the StringApp and the TISSUES database (Palasca et al., 2018) as enrichment gene set categories. Tissues sets with a gene set size below 10,000 and a false discovery rate (FDR) below 0.05 were considered significantly enriched (29 hits). These 29 tissue sets were mounted on an enrichment map using the EnrichmentMap app with a similarity cutoff of 0.5. For each of these 29 tissues set, an *Enrichment Score* was calculated as follows: first, the -log_10_ of their enrichment FDR value was plotted as a function of their gene set size. The generated dots distributed along a linear regression line (R² = 0.87) of equation (1) that we used as a baseline to calculate an *Enrichment Score* as depicted in equation (2). Tissue sets with a positive score were considered the most highly representative of the 14,646 identified mRNA set.

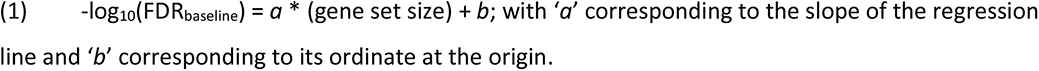

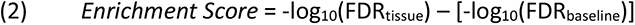

#### Unranked/ranked gene set enrichment analysis

Unranked gene set enrichment analysis (GSEA) was done using the functional enrichment tool of the StringApp and ‘*GO biological process*’, ‘*GO molecular function*’, ‘*Reactome pathways*’ and ‘*KEGG Pathways*’ as enrichment gene set categories. Ranked GSEA was done using the *Proteins with Values/Ranks - Functional Enrichment Analysis* tool of the https://string-db.org/ website. ‘*GO biological process*’, ‘*GO cellular component*’, ‘*GO molecular function*’ and ‘*KEGG Pathways*’ were used as enrichment gene set categories. FDR value below 0.05 was considered as a significant enrichment.

#### Pol_REG_ hierarchical clustering

The 166 up-regulated Pol_REG_ mRNAs were clustered based on their relative proportion in Pol, Mono and Free using the hierarchical cluster algorithm of ClusterMaker2 with pairwise average linkage and Euclidean distance metric.

#### Protein length

Protein length of the 14,646 corresponding genes were extracted from the stringdb sequence parameter of the StringApp.

#### Venn diagram

Venn diagram analyses were done using https://bioinformatics.psb.ugent.be/webtools/Venn/.

## Results

### Quality control of Sertoli cell polysome profiling

FSH-dependent molecular changes in 19-day old rat Sertoli cells were analyzed by polysome profiling, following the experimental workflow depicted in **Figure 1A**. The polysome profiles related to these samples were published previously (Musnier et al., 2012). According to their optical density profiles, Free, Mono and Pol fractions were pooled and the mRNAs in each pool were subsequently identified by next generation RNA sequencing (NGS). Of the 17,929 detected mRNAs, 14,646 genes were identified by STRING (Szklarczyk et al., 2023) (**Supplementary Table S3**). All of them but one (*Capn1*) were found in Free, Mono and Pol. Despite the variability inherent to primary cultures of cells, principal component analysis of Free, Mono and Pol replicates with or without FSH treatment grouped biological replicates together, indicating a good data reproducibility (**Figure 1B**). The pool identity accounted for most of the variance (PC1, 90%), with the Mono group found between Free and Pol as expected from its intermediate ribosomal loading status. While explaining a small portion of the variance (3%), the FSH treatment robustly explained the second principal component (PC2) of the dataset. Interestingly, Free was almost unaffected by FSH when compared to Mono and Pol which distributed similarly.

**FIGURE 1:**
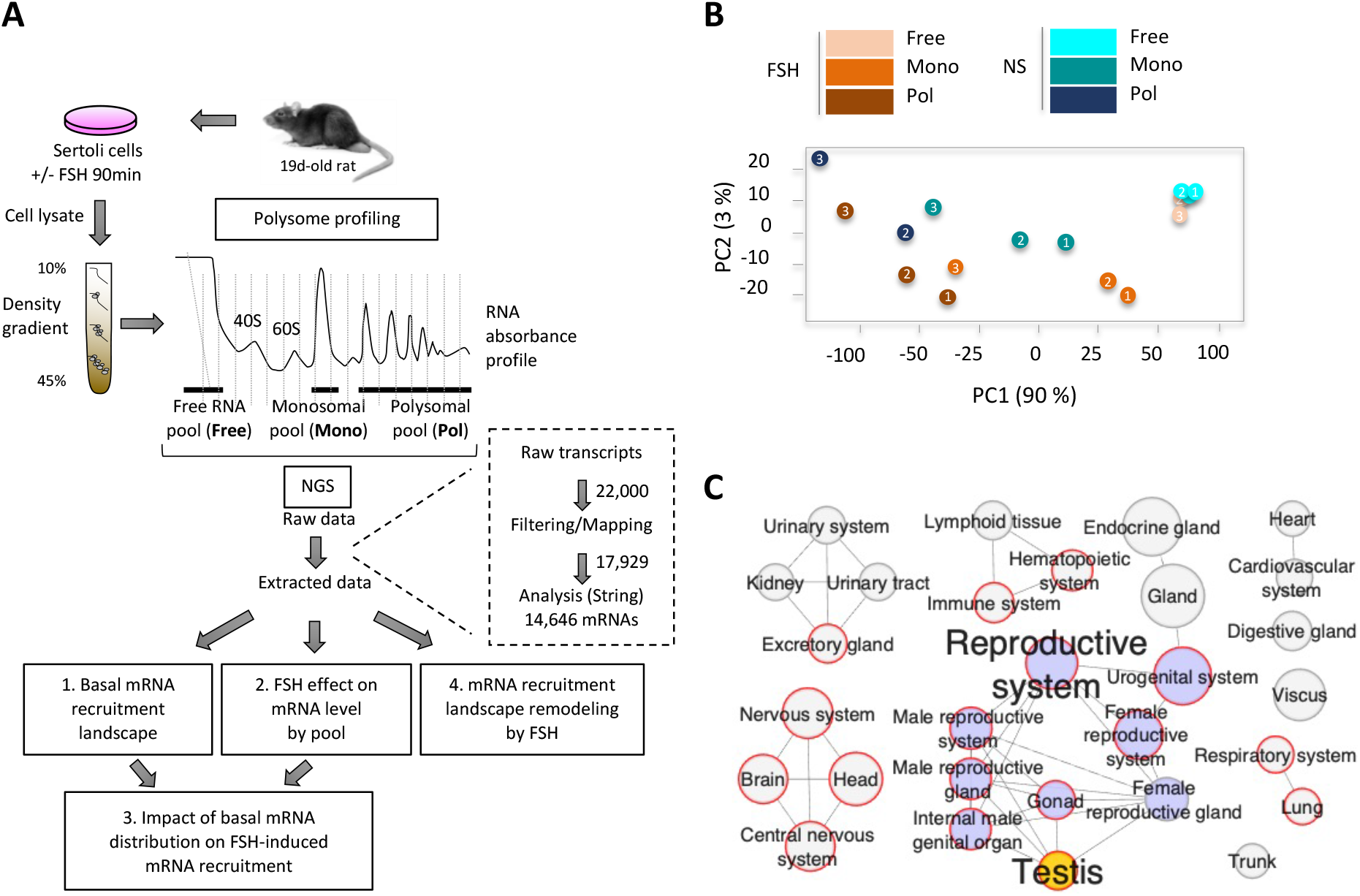
**Experimental workflow and quality control of biological samples**. (**A**) Sertoli cells from 19-day-old rats were stimulated for 90 minutes with FSH or left untreated. The cell lysates were then fractionated for polysome profiling, to collect the free (Free), monosomal (Mono) and polysomal (Pol) fractions of mRNAs. By RNA sequencing of these 3 pools exposed or not to FSH and subsequent comparison of their mRNAs content, we sequentially addressed the 4 questions depicted in the bottom part of the scheme and detailed in the text. (**B**) Principal component analysis based on Free, Mono and Pol raw mRNA counts. Dot numbers correspond to biological replicates. (**C**) Tissue gene set enrichment analysis of the 14,646 identified mRNAs. This set was compared to the rat genome from the StringApp and using the ontologies from the TISSUES database. Only tissues significantly enriched for these 14,646 genes (FDR < 0.05) and containing less than 10,000 genes are displayed. Edges represent similarity between gene sets (similarity cutoff = 0.5). Node size is proportional to gene number (from 678 for *Trunk*, to 5,234 for *Gland*). Red circles highlight tissues with positive enrichment score (see **Figure S1**).

We also checked whether Sertoli cells maintained a gene expression pattern concordant to their cell phenotype, when cultured (**Figure 1C**). Gene set enrichment analysis of the 14,646 genes using the TISSUES database as ontologies (Palasca et al., 2018) and the rat genome as reference matched testis gene expression pattern. However, many other tissues passed the 0.05 FDR threshold value. Since this statistical test is highly sensitive to the gene set size, favoring the larger sets (**Supplementary Figure S1**), we developed a simple mathematical procedure to limit this size effect and evaluated its ability to increase tissue gene set enrichment stringency, using published RNAseq data from five rat tissues (Merkin et al., 2012). This procedure confirmed the testis-specific gene expression pattern in our cultured Sertoli cells (**Supplementary Figure S1**). Among those genes, the *Inha* or *Amh* gene products are exclusively expressed in Sertoli cells (**Supplementary Table S4**). *Amh*, along with its regulator *Sox9*, encode important effectors of Sertoli cell fate. In addition, in the Human Protein Atlas, many mRNAs of this testis-specific subset are mainly expressed in Sertoli cells, when compared to other cell types of the testis. This is for example the case of *Gata4*, involved in the specific metabolic function of Sertoli cells, since it regulates the expression of genes involved in lactate metabolism. Sertoli cell-produced lactate is the energy source of spermatocytes and spermatids, instead of glucose (Boussouar and Benahmed, 2004). Another crucial function of Sertoli cells is the constitution of the blood-testis barrier (BTB), that creates a unique environment for each spermatogenetic stage. Several genes involved in the BTB formation or integrity maintenance were expressed in Sertoli cells more than in any other cell type of the testis, like *Cldn11*, *Daam2*, *Gja3*, *Nxf3* or *Serpina5*. Notably, tissues related to the nervous system also matched our mRNA dataset (**Figure 1C and Supplementary Figure S2D**). This observation is in line with the literature reporting numerous properties shared by neurons and Sertoli cells (Matos et al., 2021). In addition, the presence of terms related to the immune system is relevant to the immune surveillance achieved by Sertoli cells in the testis (Dal Secco et al., 2008; Hayrabedyan et al., 2016; Michailidis et al., 2014).

### A substantial fraction of Sertoli cell mRNAs is highly enriched in polysomes

To compare the relative enrichment of each individual mRNA in Free, Mono and Pol, the distribution of mRNAs in these 3 fractions in non-stimulated (NS) condition was equalized using DESeq2 (**Figure 2A and Supplementary Table S5**). After normalization, most mRNA distributed similarly between Free and Pol (**Figure 2B and 2C**, black arrow), between Free and Mono and between Pol and Mono (**Supplementary Figure S2A**). These results indicate that most transcripts are not randomly distributed in these three pools, and suggest that the mRNA expression level mainly dictates the content of each mRNA in each pool. However, when Pol was plotted against Free (**Figure 2B)** or Mono (**Supplementary Figure S2A**), we noticed an mRNA subpopulation with higher quantity in Pol compared to the other two pools, thus corresponding to mRNAs enriched in the polysomal pool (Pol^+^ population) (**Figure 2B and 2C**, grey arrow). Using standard DESeq2 differential expression analysis, we confirmed that the mRNAs present in this subpopulation were statistically more abundant in Pol, when compared to Mono or Free. They accounted for nearly 15% of the total mRNA population (2,084 *versus* 14,646) (**Figures 2D**, **2E and Supplementary Figure S2B**). We replicated this procedure to detect free- enriched (Free^+^) and mono-enriched (Mono^+^) subpopulations (**Figures 2D and 2E**). The Mono^+^ subpopulation was drastically smaller when compared to Pol^+^ and Free^+^, an effect attributable to its co-enrichment in mRNAs shared with Pol or Free (**Figures 2D**, 1,480 and 344 mRNAs, respectively). We also isolated free-, monosome- and polysome-depleted subpopulations (Free^-^, Mono^-^, Pol^-^respectively) as mRNAs depleted in one fraction compared to the two others, such as the 344 mRNAs depleted in Pol accounting for about 2% of the total mRNA population (**Figure 2D**). Only 83 out of 344 Pol^-^ mRNAs were shared with the 1,446 Free^+^ mRNAs, indicating that the use of Mono had a strong impact on the determination of these enriched and depleted mRNA subpopulations. Gene set enrichment analysis (GSEA) applied to all these enriched or depleted subpopulations did not detect any specific gene set ontologies. Thus, these subpopulations are not dedicated to a specialized biological function, but rather reflect the global activity of Sertoli cell. mRNAs that are the most represented in the Pol^+^ population encode proteins participating to ribonucleoprotein complex (e.g., proteins involved in splicing such as GEMIN5, SRPK2, RBM5, or in protein synthesis such as ribosomal subunit components). The Pol^-^ population contained mRNAs mainly involved in the regulation of carbohydrate metabolic processes, such as *Ppp1r3b*, *C1qtnf2*, *Adipoq* or *mTOR*. Interestingly, by comparing mRNA quantities between pools, we noticed that the Pol^+^ population contained preferentially mRNAs expressed in high amount (coordinates of the yellow cloud in **Figure 2E** and in **Supplementary Figure S2A**). Mode analysis of pools and their subpopulations confirmed the Pol^+^-specific high-expression profile, with its count-per-transcript distribution shifted toward higher values when compared to the other mRNA populations (**Figure 2F**, yellow arrow), suggesting that mRNAs that are the most expressed are also the most translated. These results are in line with those obtained in yeast, rat neurons and bacteria (Biever et al., 2020; Heyer and Moore, 2016; Lackner et al., 2007; Nguyen et al., 2022; Picard et al., 2012). As expected (Biever et al., 2020; Heyer and Moore, 2016; Picard et al., 2012), Pol^+^ mRNAs also encoded proteins with longer amino acid sequence than non Pol^+^ mRNAs (**Supplementary Figure S2C**). To calculate the relative mRNA enrichment level of each mRNA per pool, normalized mRNA values were divided by the sum of Pol, Mono and Free values. Triplot distribution confirmed the presence of a singular and substantial Pol^+^ subpopulation (**Figure 2G**). With the same procedure, the ranked GSEA based on the polysomal enrichment level applied to the 14,646 genes highlighted either a strong enrichment or a strong depletion in polysomes for mRNAs encoding proteins involved in olfactory transduction (**Figure 2H and Supplementary Figure S2D**).

**FIGURE 2:**
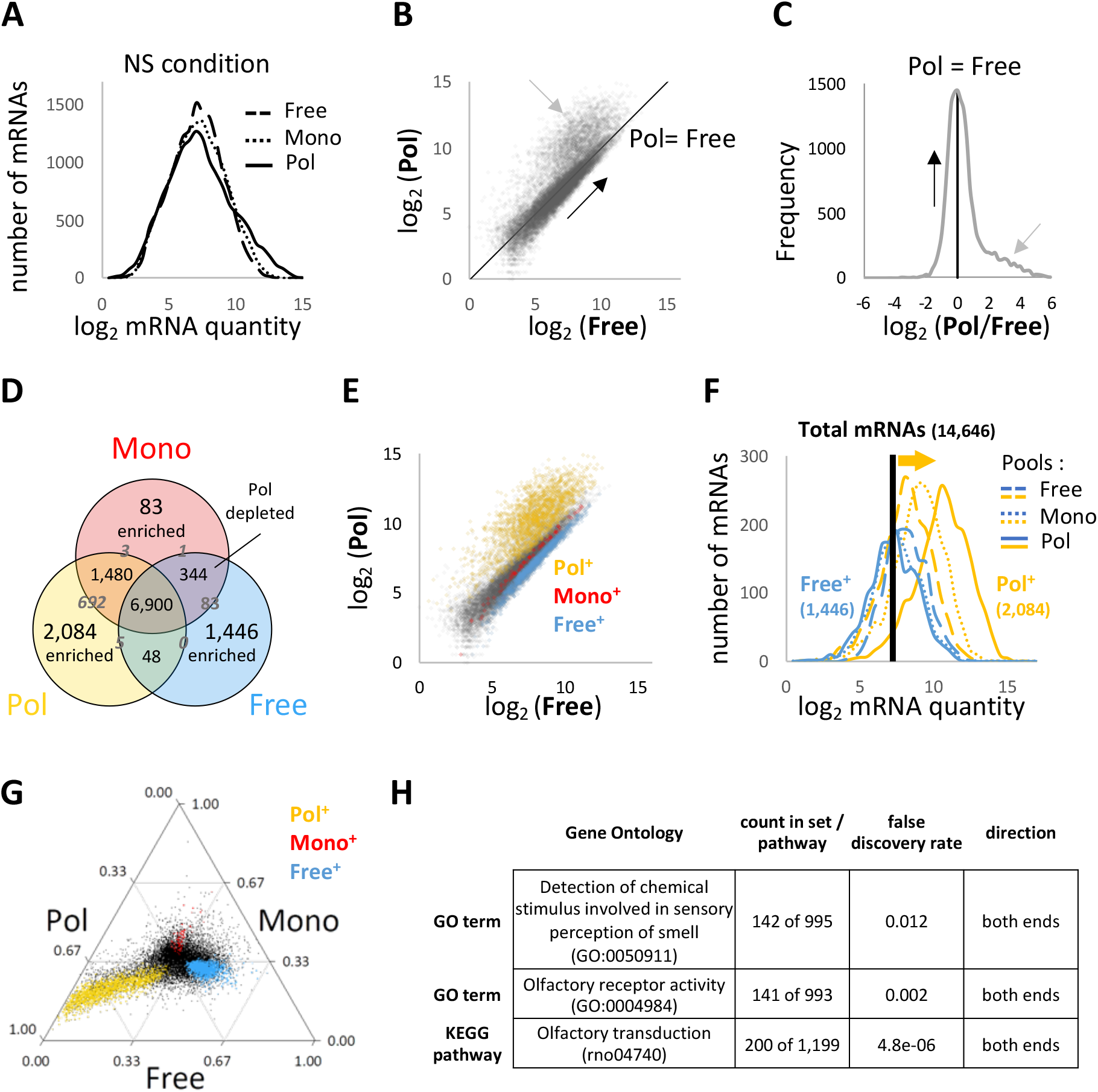
**Differences in mRNA distribution between Free, Mono and Pol**. (**A**) Normalization strategy. Free, Mono and Pol mRNA counts were normalized together by Deseq2 to maximize an even distribution of the mRNAs between the 3 pools. This normalization tends to adjust the most commonly found Pol:Free, Pol:Mono and Mono:Free ratio values to 1. (**B**) Comparison of normalized mRNAs counts in Free and Pol. Most mRNAs distributed along the diagonal (1:1 ratio, black arrow, R²= 0.71), while there is a substantial population of highly expressed mRNAs that is enriched in Pol when compared with the main distribution pattern (grey arrow). (**C**) mRNAs enrichment deviations in favor of Pol or Free compared to the main distribution pattern. Black and grey arrows represent the same populations as in (B). (**D**) Size of each mRNA set either enriched or depleted in Free, Mono or Pol, with 6,900 mRNAs being shared by the 3 pools. Enriched populations are sets of mRNAs that are statistically more represented in one pool compared to the two others. Depleted populations are sets of mRNAs that are statistically less represented in one pool compared to the two others. Grey numbers located in-between two sets indicate the number of mRNAs shared by these two sets. (**E**) Same as (B) but color-coded for enriched populations based on (D). (**F**) Comparison of the distributions of normalized mRNAs counts in Free (dashed lines), Mono (dotted lines), and Pol (plain lines) of Free^+^ (blue) and Pol^+^ (gold) mRNA populations. The mode values for the count distributions of the total mRNA population (14,646 mRNAs) in Free, Pol and Mono are similar and appear as a single black vertical bar. While the mode values for Free^+^ are similar to those for the total mRNA population, those for Pol^+^ are shifted toward higher quantity values (yellow arrow). Monosome enriched population was not plotted as its small size precludes interpretable results (83 mRNAs; data not shown). (**G**) Triplot representation of mRNAs distributions in Free, Mono and Pol after normalization. The main mRNA distribution pattern tends to distribute around a (0.33; 0.33; 0.33) coordinate at the center of the triplot, while there is a substantial polysomal-enriched mRNA subpopulation that points toward the polysomal vertex (yellow dots). (**H**) Ranked gene set enrichment analysis results based on the polysomal proportion value used for the triplot.

### Dynamics of mRNA variations in Pol, Mono and Free following FSH stimulation

The effect of FSH on the Sertoli cell translatome was estimated following standard DESeq2 normalization and differential expression method between mRNAs within the same pool (**Figure 3A**). First, mRNAs fold-change upon FSH stimulation was examined in each of the three pools (**Figure 3B and Supplementary Table S6**). In agreement with the results of the principal component analysis applied to Free, Mono and Pol replicates with or without FSH treatment (**Figure 1B**), the violin plot distribution of Free was the most compact of the three pools, while Pol was the most affected pool by the FSH stimulation. In addition, Free contained a low number of differentially regulated mRNAs (Free_REG_) with 44 and 14 mRNAs up-regulated (Free_UP_) and down-regulated (Free_DOWN_) respectively, compared to 166 (Pol_UP_) and 151 (Pol_DOWN_) differentially regulated mRNAs in Pol (Pol_REG_), and 180 (Mono_UP_) and 217 (Mono_DOWN_) in Mono (Mono_REG_) (**Figure 3C**). Twelve mRNAs were regulated in all the three pools, with two of them, *Zfp438* and *Pou2f2*, corresponding to transcription factors, and 5 of them being involved in protein processing (*Fbx30*, *Znf598*, *Rnf170*, *Dnajb2*, *TysD1*) (**Figure 3D**). All of these were up-regulated, reflecting a putative increase in their expression level upon FSH stimulation. About a third of mRNAs regulated in Pol was also found regulated in Mono (96 out of 317) and positively covariated between the two pools (**Figures 3D**). Here again, this could reflect an immediate early gene expression. Supporting this idea, the 84 mRNAs that positively covariated between Mono and Pol also had a tendency to positively covariate with Free, while the 218 mRNAs only regulated in Pol neither covariated with Mono nor with Free (**Supplementary Figure S3**). Thus, mRNAs that are the most up- or down-regulated in Pol are not the most down- or up-regulated in Mono respectively. This suggests that a principle of communicating vessel between Free, Mono and Pol does not govern the variations in FSH-induced mRNA quantity at individual mRNA level in these three pools.

**FIGURE 3:**
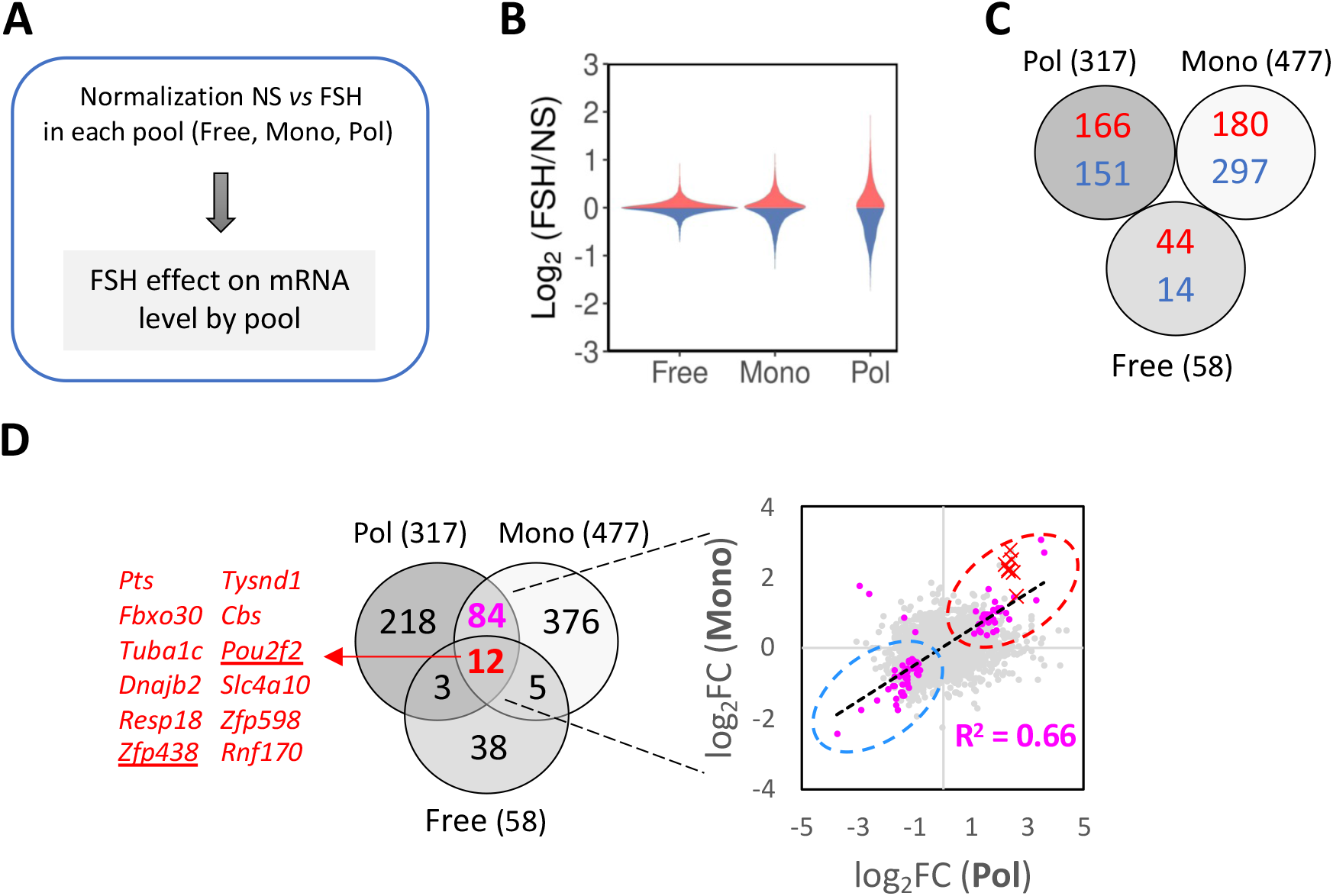
**Analysis of FSH-regulated mRNAs in the different pools**. (**A**) Normalization strategy. For each of the 3 pools (Free, Mono, Pol), NS and FSH mRNA counts were normalized together by Deseq2 to analyze the FSH effect on mRNA level by pool. (**B**) Violin plot representation of positive (red) or negative (blue) differentially regulated mRNAs in each pool. (**C**) Number of positive (red) or negative (blue) differentially regulated mRNAs in each pool. (**D**) Overlap of the differentially regulated mRNAs between Pol, Mono and Free. Eighty-four mRNAs are regulated in Pol and Mono but not in Free (pink dots). Twelve mRNAs are up-regulated in the 3 pools (red crosses). Ninety-six mRNAs (84 + 12) are regulated both in Pol and Mono. Most of them are co-recruited (45, red dotted circle) or co-depleted (47, blue dotted circle).

#### Gene ontology analysis of FSH-regulated genes

The function of the FSH-regulated mRNAs was examined by GSEA. Ranked GSEA detected enrichment for terms related to olfactory transduction in each of the three pools (**Figure 4A**). Pool enrichment or depletion for these mRNAs did not correlate between pools (**Supplementary Figure S4A**). Mono also contained statistically enriched mRNAs family sets related to ribosome components, as expected from previous works (Heyer and Moore, 2016), while Free contained one family set related to the KEGG *Estrogen signaling pathway* (data not shown). We also conducted an analysis of mRNAs regulated in Pol using functional association network in Cytoscape and the StringApp (**Figure 4B and Supplementary Figure S4B**). The top functional associations contained mRNAs related not only to olfactory transduction as in the ranked analysis, but also to translation, to vesicle-mediated transport, and to N-Glycosylation (**Figure 4B** and **Supplementary** Figure 4B). Each of these biological processes contained a ∼ 50:50 ratio of up-regulated and down-regulated mRNAs. For example, when considering mRNAs related to translation, 6 of them (*Larp1*, *Mrpl16*, *Mrps34*, *Rpl18a*, *Rpl24*, *Rps19*) were down-regulated while 4 (*Cars*, *Eif5*, *Rpl41*, *Zfp598*) were up-regulated. This may reflect the action of FSH on negative regulator of 5’TOP mRNA such as *Larp1*, which is consistent with the regulatory role of FSH on mRNAs having this motif (Tréfier et al., 2018b).

**Figure 4:**
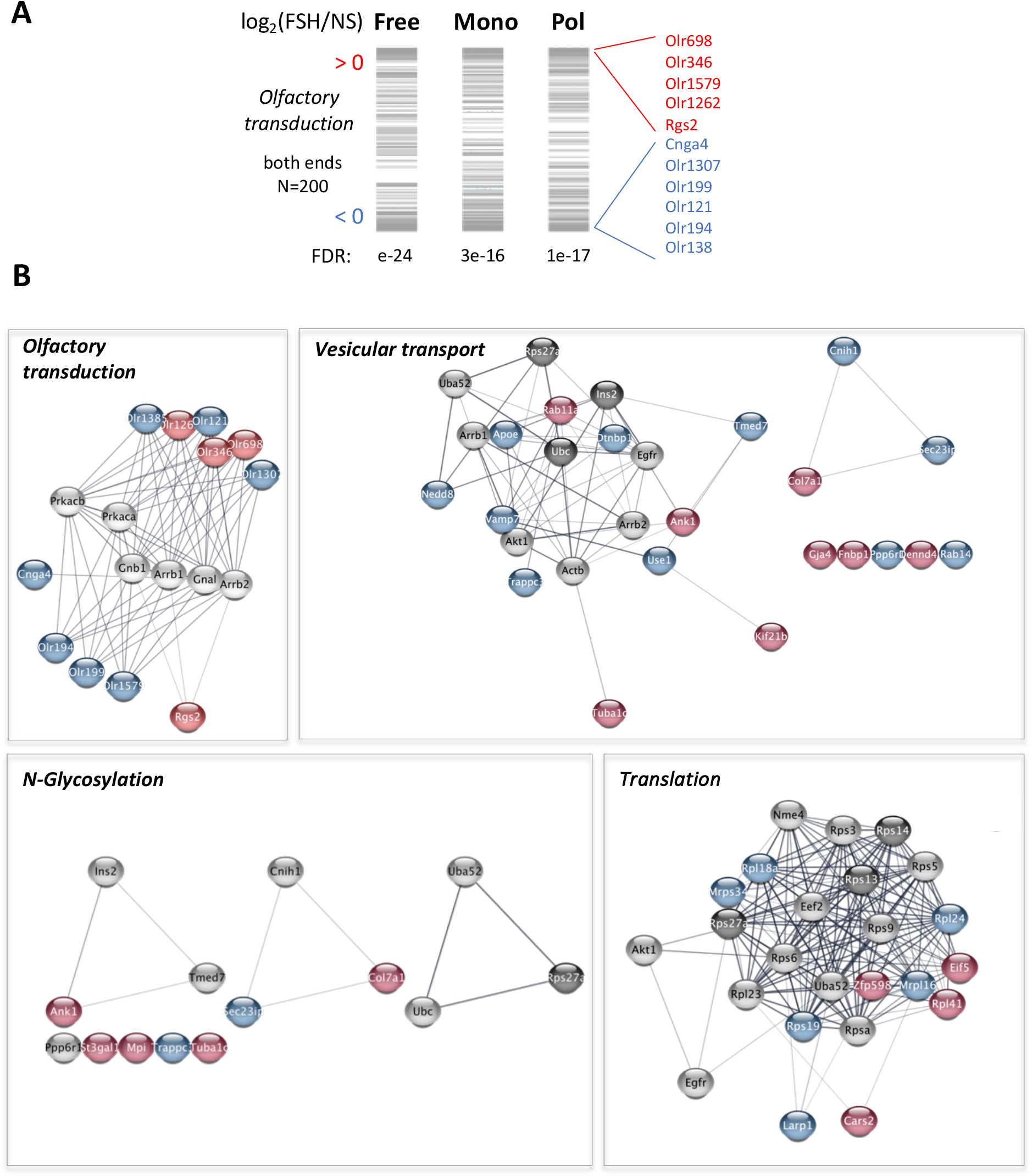
**Gene ontology analysis of FSH-regulated genes**. (**A**) Ranked gene set enrichment analysis position profiles based on FSH/NS fold change in Free, Mono and Pol. Grey bars indicate the rank of the 200 mRNAs of the KEGG olfactory transduction gene set: the upper value in the profile is the mRNA with the highest log_2_(FSH/NS) value, and conversely. The names of the statistically up-regulated (red) and down-regulated (blue) mRNAs in Pol are indicated. (**B**) Main functional association networks of the 317 Pol_REG_ mRNAs added with additional 10% mRNAs from the background, with FDR<0.05. In red, mRNA up-regulated in Pol; in blue, mRNA down-regulated in Pol; in pale grey, mRNA of the data set that are not regulated by FSH; in dark grey, added nodes that are not present in the dataset. The networks were generated in Cytoscape with the StringApp using a confidence score of 0.4.

Hence, these data suggest that 90 min FSH exposure induced a strong reorganization of olfactory receptor expression pattern as well as other basic biological functions. Since mRNAs related to olfactory transduction were also highly segregated between Free, Mono and Pol (**Supplementary Figure S2D**), this raises the intriguing possibility that the initial polysomal enrichment level (i.e. prior to FSH stimulation) strongly impacts FSH effect.

### Polysomal enrichment level before stimulation is a major determinant of short term, FSH-induced polysomal recruitment

To assess whether the basal mRNA distribution impacts FSH-induced mRNA recruitment to polysomes, we first confronted the FSH effect on the polysomal content to the mRNA enrichment level in polysomes before stimulation (**Figure 5A**). Pol_DOWN_ were 3.6 times more enriched in Pol^+^ when compared to basal mRNA distribution, with half (52 %) of Pol_DOWN_ originating from Pol^+^ (**Figure 5B**). Conversely, Pol_UP_ were higly enriched (7.4x) in mRNAs originating from Pol^-^. Plotting the FSH-induced fold change of counts in Pol as a function of the basal polysomal enrichment confirmed the strong segregation between up-regulated and down-regulated Pol_REG_ (**Figure 5C**): up-regulated mRNAs corresponded to mRNAs that were depleted from polysomes before FSH stimulation (low % Pol NS), while down-regulated mRNAs corresponded to mRNAs that were enriched in polysomes in basal conditions (high % Pol NS). This indicates that the distributions of mRNAs enriched or depleted in the polysomes are out of equilibrium at the basal level, and tend to shift toward a more balanced distribution following FSH stimulation: polysome enriched mRNAs tend to be released from polysomes upon FSH stimulation, while polysome depleted mRNA tend to be recruited.

**FIGURE 5:**
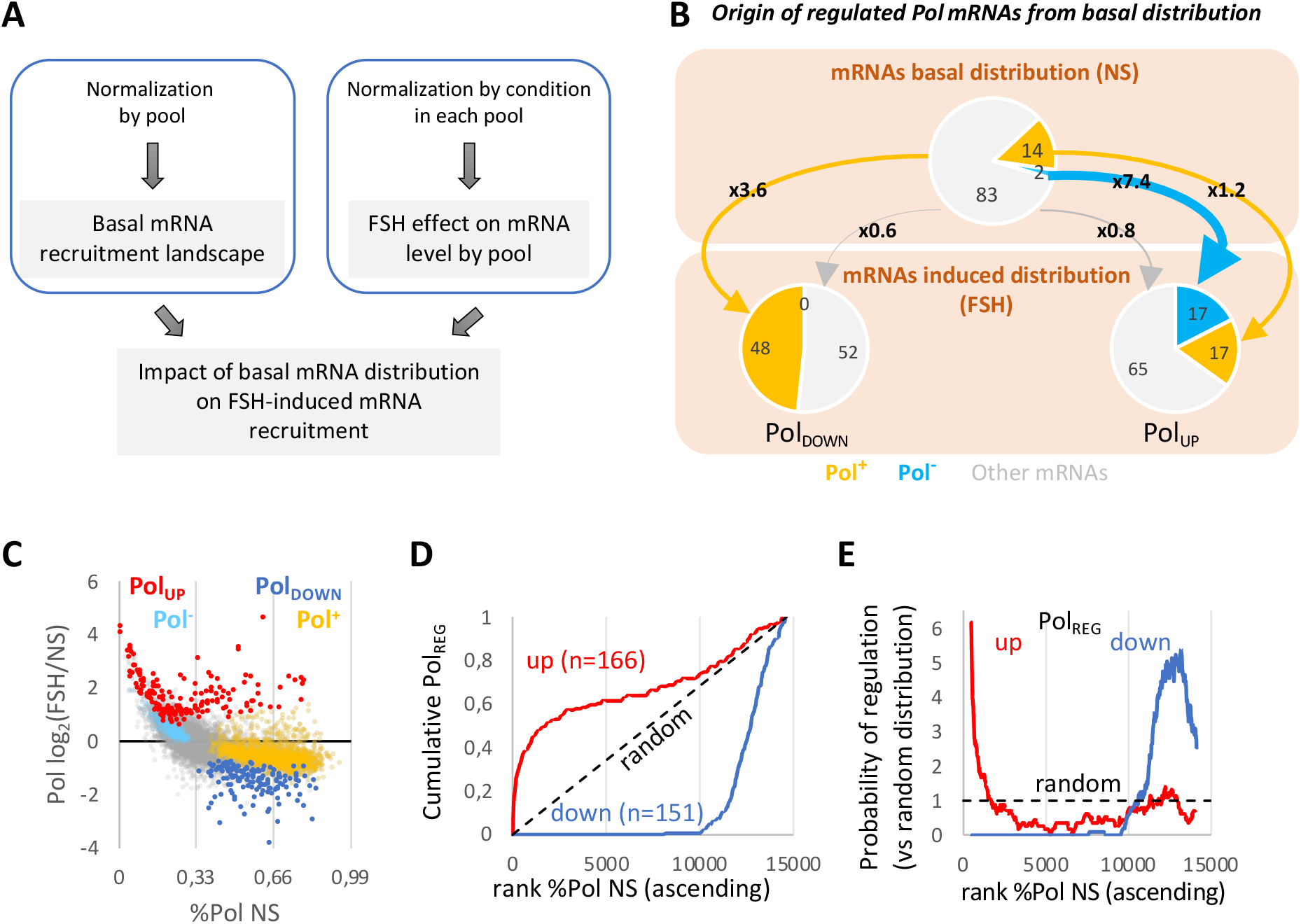
Impact of the basal mRNA distribution on FSH-induced mRNA recruitment to polysomes. (**A**) Analysis strategy. The effect of FSH on the mRNA level in each pool was confronted with the mRNA distribution landscape before FSH stimulation. (**B**) Origin of regulated Pol mRNAs from the basal distribution. Forty-eight % of Pol_DOWN_ mRNAs originate from Pol^+^, which represents a 3.6-fold enrichment in Pol^+^ compared to the total mRNA population (48%/14% = 3.6). (**C**) Plot of mRNAs counts fold-change in Pol_REG_ as a function of their basal enrichment levels in Pol (% Pol NS). Red and blue populations represent FSH up– and down-regulated mRNAs, respectively. Gold and cyan populations represent Pol^+^ and Pol^-^ in non-stimulated cells, respectively. Note that red and cyan populations overlap and locate on the upper-left corner of the graph, while blue and gold populations overlap and locate on the lower-right corner of the graph. (**D**) Cumulative distributions of up- and down-Pol_REG_ based on mRNA enrichment level in Pol before FSH stimulation (rank % Pol NS). The 14,646 mRNAs were ranked in an ascending order from the lowest % Pol NS to the highest % Pol NS. (**E**) Probability for an mRNA to be up- or down-regulated in Pol, based on its enrichment level in Pol before FSH stimulation (%Pol NS). The probability of regulation is calculated using a moving average of 1,000 genes.

Next, we analyzed this effect at the scale of the whole Sertoli cell mRNA population using cumulative distribution of either up or down Pol_REG_ as a function of their polysomal enrichment level before stimulation (**Figure 5D**). This confirmed that the FSH-induced polysomal recruitment was extremely sensitive to the basal polysomal recruitment in an oriented and consistent manner: mRNAs that were already enriched in polysomes were more released, while those that were already depleted from polysomes were more recruited upon hormonal input (**Figure 5E**). In particular, 99% of down- regulated mRNAs came from the top third most polysomal-enriched mRNAs prior to stimulation (**Figure 5D**, ranks 10,000 and over out of 14,646). For upregulated mRNAs, although 60% of them came from the top third most polysomal-depleted mRNAs prior to stimulation (**Figure 5D**, ranks 0 to 5,000), 25% of them came from the top third most polysomal-enriched mRNAs (**Figure 5D**, ranks 10,000 and over out of 14,646). Trying to understand the origin of these 25% “escapers” (**Supplementary Table S7**), we applied a hierarchical clustering to the up Pol_REG_ based on their enrichment level in each pool before stimulation (**Supplementary Figure S5A**). Strikingly, these 25% escapers were also mostly up- regulated in Mono and included all the mRNAs that were statistically up-regulated in the three pools (**Figure 3D**). Expressing the cumulative distribution of Pol_REG_ with mRNAs regulated only in Pol corrected most of the asymmetry observed between up and down Pol_REG_ curves (**Supplementary Figure S5B**). Thus, a tempting explanation could be that these escapers actually correspond to mRNAs with FSH-induced increased transcription, rather than pure polysomal recruitment at constant mRNA expression level.

### FSH-induced mRNA transfer between pools does not affect the global Sertoli cell mRNA distribution landscape

Our previous results on the mRNA distribution between Free, Mono and Pol in resting Sertoli cells deciphered a peculiar distribution landscape with one main distribution pattern and a prominent polysomal enriched mRNA subpopulation (**Figure 2G**). We wondered whether FSH treatment would alter this distribution pattern. Thus, we normalized once more Free, Mono and Pol using DESeq2, but this time in pools obtained after FSH treatment (**Figures 6A**). The mRNA distribution pattern once again exhibited the prominent polysomal enriched subpopulation and remained mostly unaffected (**Figure 6B**). Nevertheless, we also noticed substantial changes, with a global decrease, rather than replacement (**Supplementary Figure S6A**), of mRNAs enriched or depleted in each pool (**Figures 6B-C and Supplementary Figure S6B**), while up and down Pol_REG_ showed extensive coordinate shifts in triplot representations (**Figure 6D**).

**FIGURE 6:**
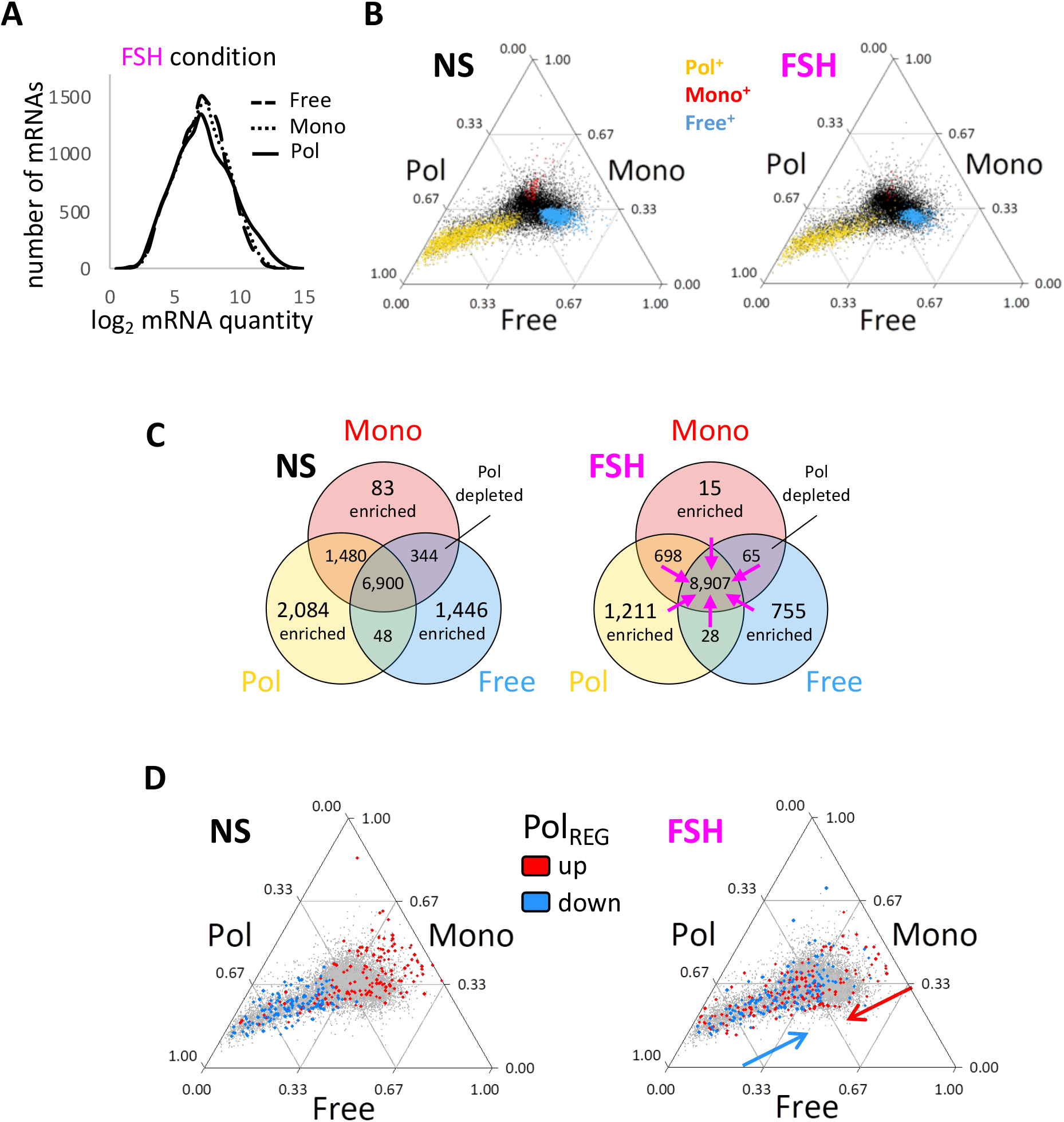
mRNA distribution landscape remodeling by FSH. (**A**) Normalization strategy. Free, Mono and Pol mRNA counts in FSH condition were normalized together by Deseq2 to maximize an even distribution of the mRNAs between the 3 pools, as what was done for the non-stimulated condition (see Figure 2A). (**B**) Triplot representation of FSH compared to basal (NS) mRNAs distributions in Free, Mono and Pol. Enriched populations are color-coded. (**C**) Size number of each mRNAs sets either enriched or depleted in Free, Mono or Pol in FSH *versus* basal (NS) conditions. Pink arrows: FSH induced a reduction of both enriched and depleted mRNA populations. (**D**) Triplot representation of Pol_REG_ distribution in FSH compared to basal (NS) conditions. The blue and red arrows indicate the global shift of the down-regulated and up-regulated mRNAs in the triplot, respectively.

Collectively, these results indicate that the FSH-induced reassignment of ribosomes to specific mRNAs has to comply with a global mRNA distribution landscape tightly maintained around a set point where most of mRNAs distribute similarly between Free, Mono and Pol, while a subset of mRNAs gets highly enriched in Pol.

## Discussion

In this study, we combined polysome profiling, RNA sequencing, and a model of mRNA distribution across three distinct ribosome-bound pools (free, monosomal, and polysomal) to characterize the FSH-regulated dynamics of mRNA translation in Sertoli cells. First, we observed that, in unstimulated cells, mRNA distribution followed a bimodal pattern: the majority of the mRNAs (85%) adhered to a standard mode, while a subset (15%) displayed a specialized mode marked by a high expression level and enrichment in polysomes. FSH stimulation induced a global reorganization of the translatome associated to Sertoli cell core biological functions, including metabolism, transcription, translation, vesicular transport, etc. Importantly, mRNA recruitment to or release from polysomes was heavily influenced by their baseline polysomal enrichment levels, while the overall mRNA distribution among the three pools remained largely stable. These findings suggest that Sertoli cells operate under constraints of limited ribosomal availability, requiring a redistribution of ribosomes from polysome-enriched to polysome-depleted mRNAs during hormonal adaptation.

Previous mathematical models have proposed that ribosomal availability limits simultaneous translation of numerous mRNAs (Jain et al., 2022; Raveh et al., 2016; Shah et al., 2013). However, this assumption has been experimentally challenged in yeast, where ribosomal proteins were found in excess and only used for translation under conditions of extreme need (Metzl-Raz et al., 2017). Nevertheless, exploring the consequences of defective synthesis of ribosomal 40S and 60S subunits due to mutations found in human ribosomopathies have reconciled competing hypotheses, including mRNA competition for ribosomes (Cheng et al., 2019) and translational selectivity (Khajuria et al., 2018). Overall, whether the number of ribosomes ready for translation is limiting or not seems to rely on the cellular type, state and stimulus.

To our knowledge, the bi-modal distribution of mRNAs assignment to polysomes has not been documented in the literature. The lack of prior documentation may stem from the fact that ribosome occupancy, i.e. the fraction of transcripts that are bound to at least one ribosome, thus including Mono, is usually chosen as a metric to characterize the mRNAs distribution to ribosomes (Arava et al., 2003; Lackner et al., 2007; Nguyen et al., 2022; Picard et al., 2012). However, monosomes are increasingly recognized as a unique pool with specific characteristics, that differ from those of the polysomes (Biever et al., 2020; Heyer and Moore, 2016), a conclusion consistent with our new observations. Hence, since Mono accounts for a substantial amount (∼14%) of the mRNA content of a cell (Arava et al., 2003), by treating monosomes and polysomes separately, we detected differences obscured when these pools are merged. Additionally, our approach compared polysomal mRNAs to the free pool, rather than the total mRNA content, revealing a broader distribution than typically reported. Since Pol contains the major fraction of an mRNA in a cell, exceeding 50% in average (Arava et al., 2003; Kawaguchi and Bailey-Serres, 2005), comparing Pol to the total mRNA content produces a more compact and homogeneous distribution than when comparing to Free as a reference. We ruled out that the distribution of mRNAs assignment to polysomes would depend on the cell type, the harvesting method to extract the cells, and the organism of interest, because ribosome occupancy is well conserved in distinct organisms and cell types.

Several publications already reported a positive correlation between the expression level of an mRNA and its ribosome occupancy (Biever et al., 2020; Heyer and Moore, 2016; Lackner et al., 2007; Nguyen et al., 2022; Picard et al., 2012). The high expression level of Pol^+^ mRNAs was reported in two of these studies which specifically compared Mono^+^ and Pol^+^ mRNAs in yeast and rat neurons (Biever et al., 2020; Heyer and Moore, 2016). In yeast, these highly expressed mRNAs encoded highly abundant proteins (Heyer and Moore, 2016). Since increasing translation over transcription minimizes the production cost per protein (Frumkin et al., 2017; Hausser et al., 2019), favoring a high translation over transcription ratio for highly abundant proteins could save a substantial and critical amount of cellular energy. Thus, Pol^+^ might originate from energy expense optimization of gene expression. However, the functional role of these mRNAs remains unclear. Our GSEA results indicate that Pol^+^ reflects the global mRNA content of the Sertoli cell. In yeast, Pol^+^ was enriched in mRNAs coding for proteins involved in translation, and general metabolism (Heyer and Moore, 2016), while in rat neurons they were linked to intracellular vesicle and more general metabolism processes (respiratory chain, proteasome, etc) (Biever et al., 2020). Further investigation is needed to clarify Pol^+^ roles in Sertoli cells.

Interestingly, our results support the new concept of a saturating level of polysomal enrichment for a given mRNA. Approximately 25% of mRNAs recruited to polysomes under FSH stimulation were already polysome-enriched before stimulation, suggesting that their increased abundance likely stemmed from transcriptional upregulation rather than enhanced recruitment to polysomes. This conclusion is supported by i/ their concurrent increase in the monosomal and polysomal pools without changes in the free pool, and ii/ they contained all the cellular mRNAs concurrently up-regulated in Free, Mono and Pol. Note that a regulation decreasing the decay of these mRNAs would also theoretically provide the same outcome (Heck and Wilusz, 2018). It can be speculated that when there is a saturating number of ribosomes relative to the number of copies of a given mRNA, the steady state copy number of this mRNA has to increase in order to increase the production of the encoded protein to adapt to a new condition. Although the existence of such backward dialog from translation to transcription has yet to be demonstrated, this positive coupling would be another mechanism, distinct from energy expense optimization of gene expression (Frumkin et al., 2017; Hausser et al., 2019), explaining why polysome-enriched mRNAs are also the most expressed. Comparison of the dynamics of transcription and translation following hormonal stimulation should provide more insight into this intriguing phenomenon.

Despite the global stability of mRNA distribution landscape among Free, Mono and Pol, FSH stimulation reduced the number of mRNAs enriched in any single pool. Since FSH is known to promote anabolic activities in Sertoli cells (Santi et al., 2020), this shrinkage of enriched or depleted mRNAs subpopulations could be a marker of such metabolic state transitions. In agreement, a symmetric situation is illustrated by plant dehydration, a stress known to decrease the cellular metabolic activity (Lawlor and Tezara, 2009) that was accompanied by an increase of the mRNA distribution heterogeneity (Kawaguchi et al., 2004).

Our prior work, that reported the polysome profiling of the samples used here (Musnier et al., 2012), supported an FSH-induced transfer of mRNAs from Mono to Pol, in agreement with the *bulk mRNA transfer* well described in other cell types and conditions. However, this bulk trend does not explain *individual mRNA dynamics* observed here, which instead suggest that mRNA synthesis and decay dominate over inter-pool redistribution. We assume that the sequential normalizations that were done here provide more precise relative quantification than the experimental quantification that was done in our previous work. For some unclear reason, the 2 mRNAs that were described as selectively translated in (Musnier et al., 2012) by qRT-PCR did not reach significance in Pol in this study. In the FSH-regulated translatome, enrichment was detected solely for mRNAs encoding olfactory receptors, which are minimally expressed in Sertoli cells relative to germ cells (Parmentier et al., 1992; Soffientini et al., 2017). Ectopic ORs have been shown to promote spermatozoa chemosensing, that might be involved in the sperm-egg dialog, both in human and in a mouse model (Fukuda et al., 2004; Spehr et al., 2003). Although their function remains unclear in Sertoli cells, ORs have been shown to interact with selective GPCRs such as the β2- adrenoceptor (Hague et al., 2004), which is expressed in Sertoli cells (Troispoux et al., 1998). This raises the possibility of regulatory interactions of ORs, affecting GPCR signaling or trafficking in Sertoli cells.

Interestingly, while Sertoli cell functions like chromatin remodeling or GPCR signaling were not enriched upon FSH stimulation, the same functions were accomplished with different effector proteins, indicative of a functional rewiring of Sertoli cell activity. Of note, Sertoli cells are epithelioid, polarized cells, that exert highly specialized functions specific of each spermatogenic stage that lie on their vertical axis. Hence, local function enrichment might be averaged when mRNAs of the whole cell are analyzed. This point is further supported by many studies in neurons, where specific mRNA translation has been detected in dendritic spines during neuronal activity (Biever et al., 2020; Bramham and Wells, 2007; Ostroff et al., 2002) In conclusion, we report the first genome-wide translatomic analysis of FSH-stimulated Sertoli cells. Our findings reveal that the basal polysomal enrichment of an mRNA dictates its propensity to be translationally responsive to hormonal signals. This highlights the importance of considering baseline mRNA distributions when interpreting changes in polysomal recruitment. Since Sertoli cells are polarized, future work should investigate whether localized translation supports the diverse functional demands of Sertoli cells across spermatogenic stages and whether it is sensitive to hormone stimulation (Tréfier et al., 2018c).

## Acknowledgements

The authors acknowledge Professor George Bousfield for the kind gift of purified porcine FSH. We thank the technicians of PAO at the rodent INRAE 420 Animal Physiology Facility (https://doi.org/10.15454/1.5573896321728955E12) for the care and breeding of Wistar rats. We acknowledge the sequencing and bioinformatics expertise of the I2BC High-throughput sequencing facility, supported by France Génomique (funded by the French National Program "Investissement d’Avenir" ANR-10-INBS-09). We acknowledge the use of ChatGPT [https://chat.openai.com/] for edition of the last draft of this paper.

## Author contribution

Conceptualization: PC, XL, JM, FG and RY Data curation: TB, HO, RC and AP Formal analysis: XL, NA, AT and RY Funding acquisition: PC, LP, ER and RY Investigation: XL, AM, KL, XL, SB, YJ, AT, LD and EP Methodology: XL, RC, HO, RY, NA, AT Project administration: PC Resources: JM Software: NA Supervision: PC and RY Validation: LD, LP and EP Visualization: XL, RY and NA Writing- original draft: XL and PC Writing- review and editing: All authors contributed significant revisions to later versions and approved the final version before submission.

## Abbreviations

BTB: blood-testis barrier
DEG: differentially expressed gene DEPC, diethylpyrocarbonate
DTT: dithiothreitol
EDTA: ethylenediaminetetraacetic acid ET1R, endothelin 1 receptor
FDR: false discovery rate Free, free pool
Free^-^: deprived in the free pool Free^+^, enriched in the free pool
Free_REG_: FSH-regulated in the free pool
Free_DOWN_: down- regulated by FSH in the free pool Free_UP_, up- regulated by FSH in the free pool
FSH: follicle-stimulating hormone
FSHR: follicle-stimulating hormone receptor GNRHR, gonadotropin-releasing hormone receptor GPCR, G protein-coupled receptor
GSEA: Gene set enrichment analysis IFN𝛾, interferon gamma
LH: luteinizing hormone
mGluR1: metabotropic glutamate receptor 1 Mono, monosomal pool
Mono^-^: deprived in the monosomal pool Mono^+^, enriched in the monosomal pool
Mono_REG_: FSH-regulated in the monosomal pool mTOR, mammalian target of rapamycin
NGS: next-generation sequencing PBS, phosphate-buffered saline Pol, polysomal pool
Pol^-^: deprived in the polysomal pool Pol^+^, enriched in the polysomal pool
Pol_REG_: FSH-regulated in the polysomal pool
Pol_DOWN_: down- regulated by FSH in the polysomal pool Pol_UP_, up- regulated by FSH in the polysomal pool
SDS: sodium dodecyl sulfate
TRAP: tandem ribosome affinity purification

