## Supplementary Material for "Hormone-regulated dynamics of mRNA distribution on ribosomes in Sertoli cells"

- Supplementary Figures S1 to S6
- Legends to Supplementary tables
- References to Table S4

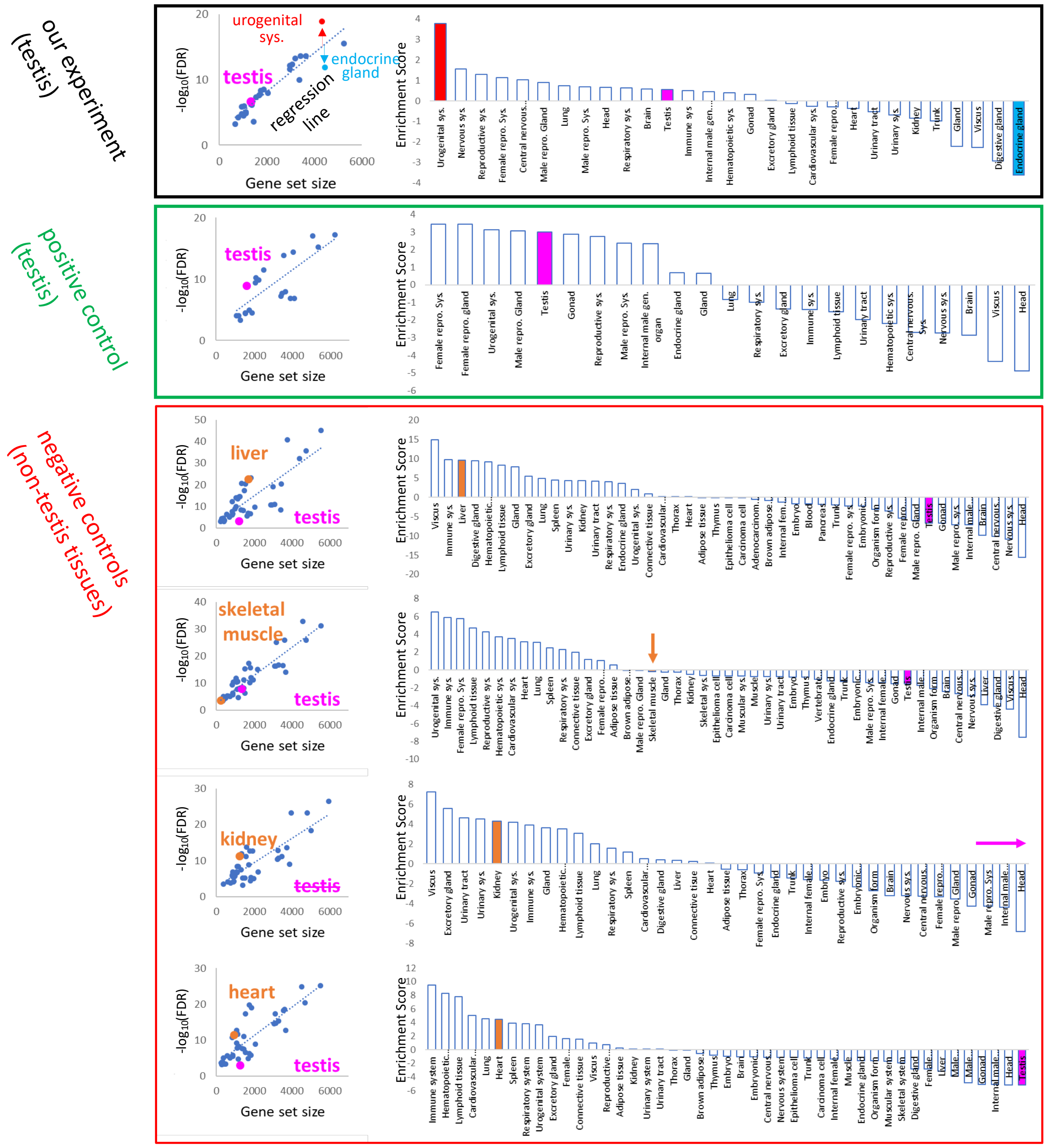


**FIGURE S1, related to Figure 1**: **Method to increase the tissue gene set enrichment analysis stringency**. **Black box**, left panel: plots of the -log$_{10}$ FDR value of the remaining 29 significantly enriched tissues (see **Figure 1C**) as a function of their gene set size. These tissues distribute along a linear regression line that is used as a baseline to calculate an Enrichment Score for each of these tissues, the higher the distance, the higher the score. This score corresponds to the difference between the -log$_{10}$ FDR value of the tissue (y-coordinate) and its corresponding -log$_{10}$ FDR value for a gene set with the same size according to the linear regression line. Thus, tissues with upward positions relative to the regression line get a positive score (red arrow) and correspond the most to the gene set tested (14,646 genes from the cultured Sertoli cells here), while tissues with downward positions relative to the regression line get a negative score (blue arrow). The purple dot indicates the testis. Right panel: histograms of the Enrichment Score of each of the 29 tissues ranked from higher to lower score. Purple color indicates testis. Red, blue, and purple histograms represent the same tissues as in the left panel. Green and Red boxes: Gene set enrichment analyses of published RNA-seq data (Merkin et al., 2012) from five rat tissues used as positive (green box; testis) and negative (red box; liver, skeletal muscle, kidney, heart) controls. The testis tissue obtained a positive score for its equivalent TISSUES gene set (green box, purple bar), while the other 4 non-testis tissues obtained a positive score for their equivalent TISSUES gene set (red box, orange bar) and a negative score for testis (purple bar), thus indicating robustness of the method.

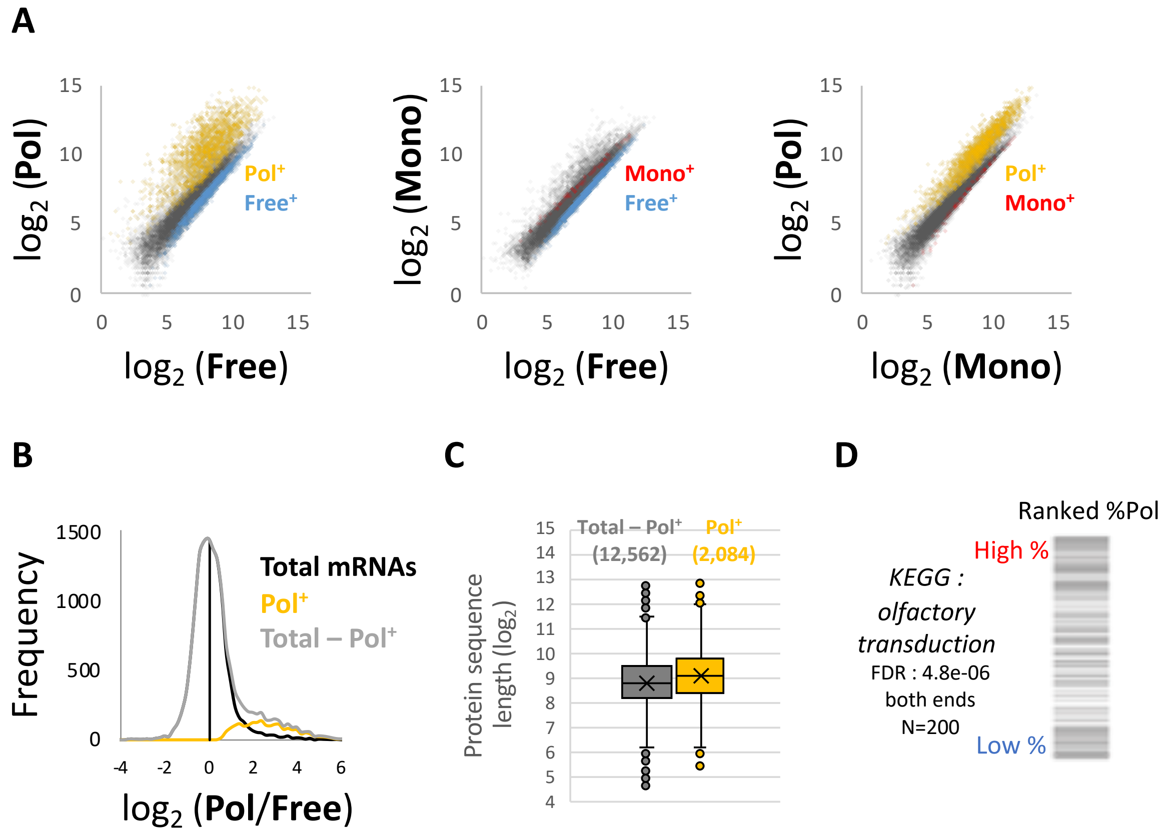


**FIGURE S2, related to Figure 2**: **Differences in mRNA distribution between Free, Mono and Pol**.
(**A)** Comparison of normalized mRNAs counts in Free and Pol (left), Free and Mono (center), Mono and Pol (right), which are color-coded for enriched populations based on Figure 2D. **(B)** mRNAs enrichment deviations in favor of Pol or Free compared to the main distribution pattern. One distribution pattern corresponds to the main distribution pattern (grey, Total – Pol$^{+}$) while the other one consists in mRNAs highly enriched in Pol (yellow, Pol$^{+}$). **(C)** Comparison of the protein sequence length encoded by the Pol$^{+}$ or non- Pol$^{+}$ mRNA populations. **(D)** Ranked gene set enrichment analysis position profile of the 14,646 genes based on % Pol. Grey bars indicate the position of the 200 mRNAs of the KEGG olfactory transduction gene set. The upper value is the mRNA the most enriched in Pol (high % Pol), while the lower value is the mRNA the most depleted in Pol (low % Pol).

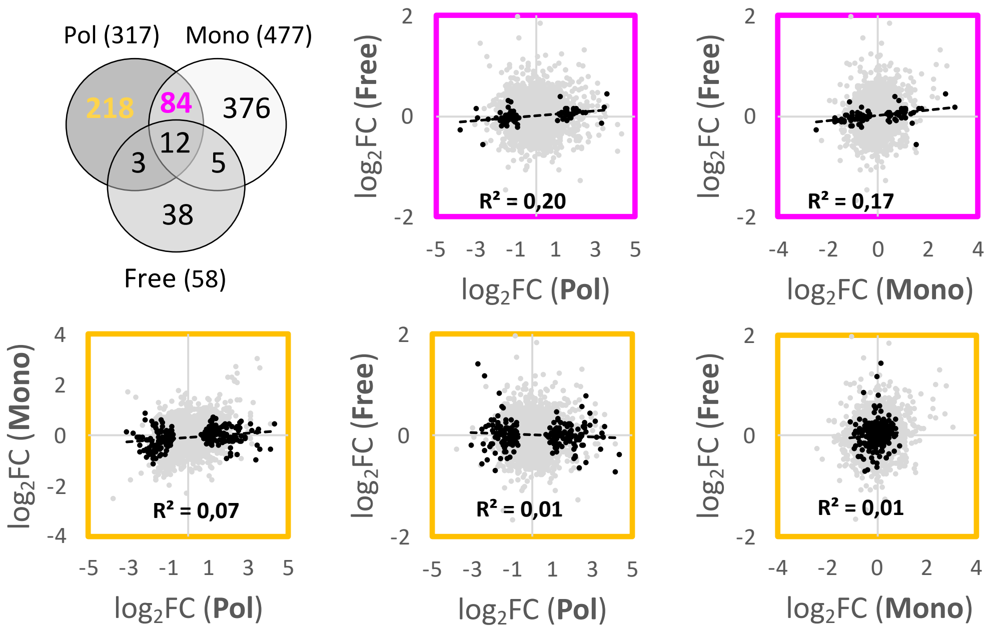


**FIGURE S3, related to Figure 3**: **Comparison of Pol, Mono and Free fold changes of FSH-regulated mRNAs**. The Venn diagram represents the overlap between mRNAs differentially regulated in Pol, Mono and Free. Two hundreds and eighteen mRNAs are only regulated in Pol. Eighty-four mRNAs are regulated in Pol and Mono but not in Free. In the pink plots, the fold changes of the 84 mRNAs in Pol and Mono are compared with the fold changes in Free. Note that they tend to positively covariate. Gold plots: Comparison of Pol, Mono and Free fold changes of the 218 mRNAs. These mRNAs do not covariate.

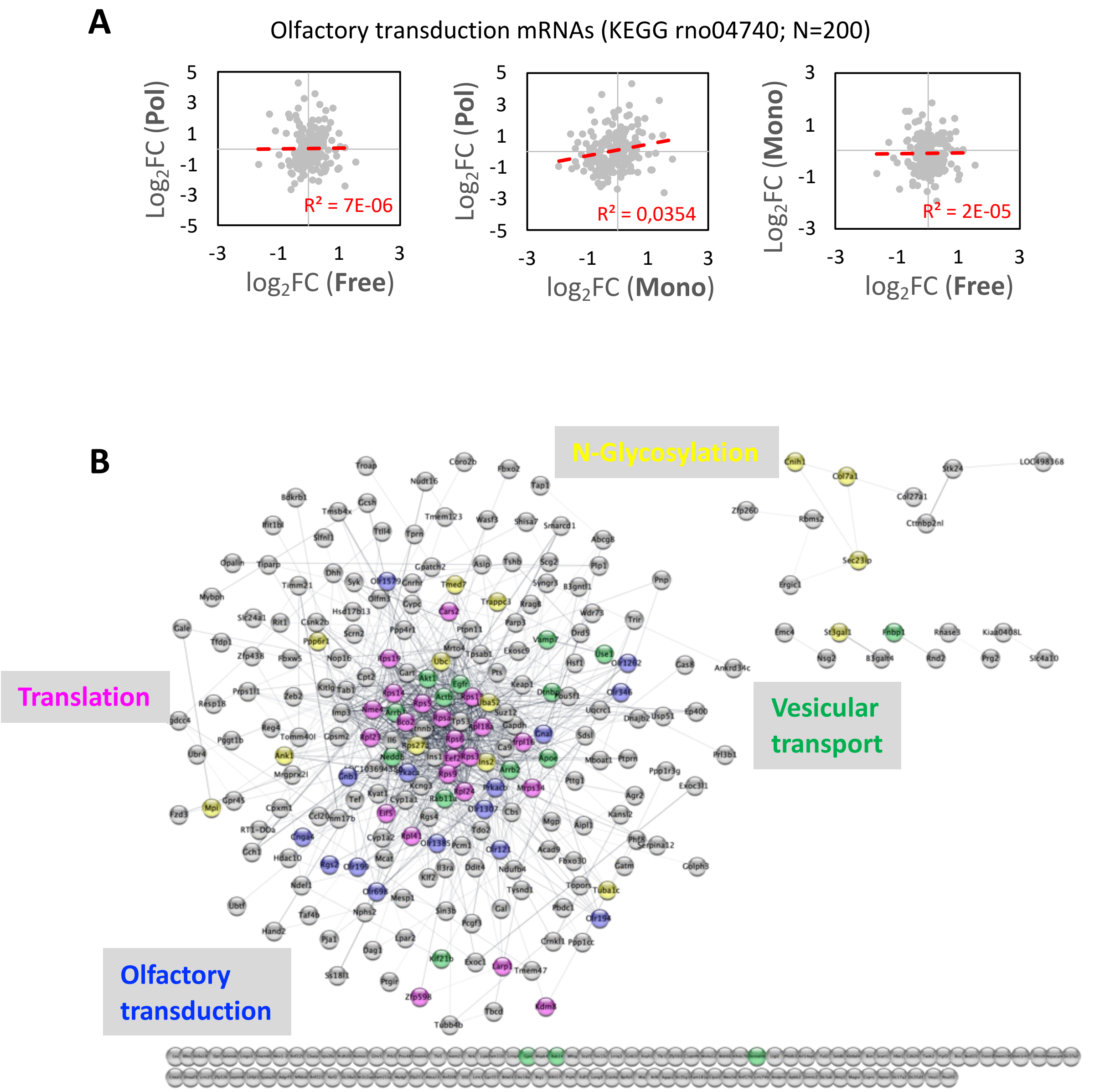


**FIGURE S4, related to Figure 4. Gene ontology analysis of FSH-regulated genes.** (**A**) Comparison of Pol, Mono and Free fold changes of the 200 mRNAs of the KEGG olfactory transduction gene set. (**B**) Functional association network of the 317 Pol differentially regulated mRNAs as in Figure 4B but with all mRNA names. The network was generated by Cytoscape and the String app using a confidence score of 0.4 and a force-directed layout. mRNAs related to the main biological ontologies are color-coded.

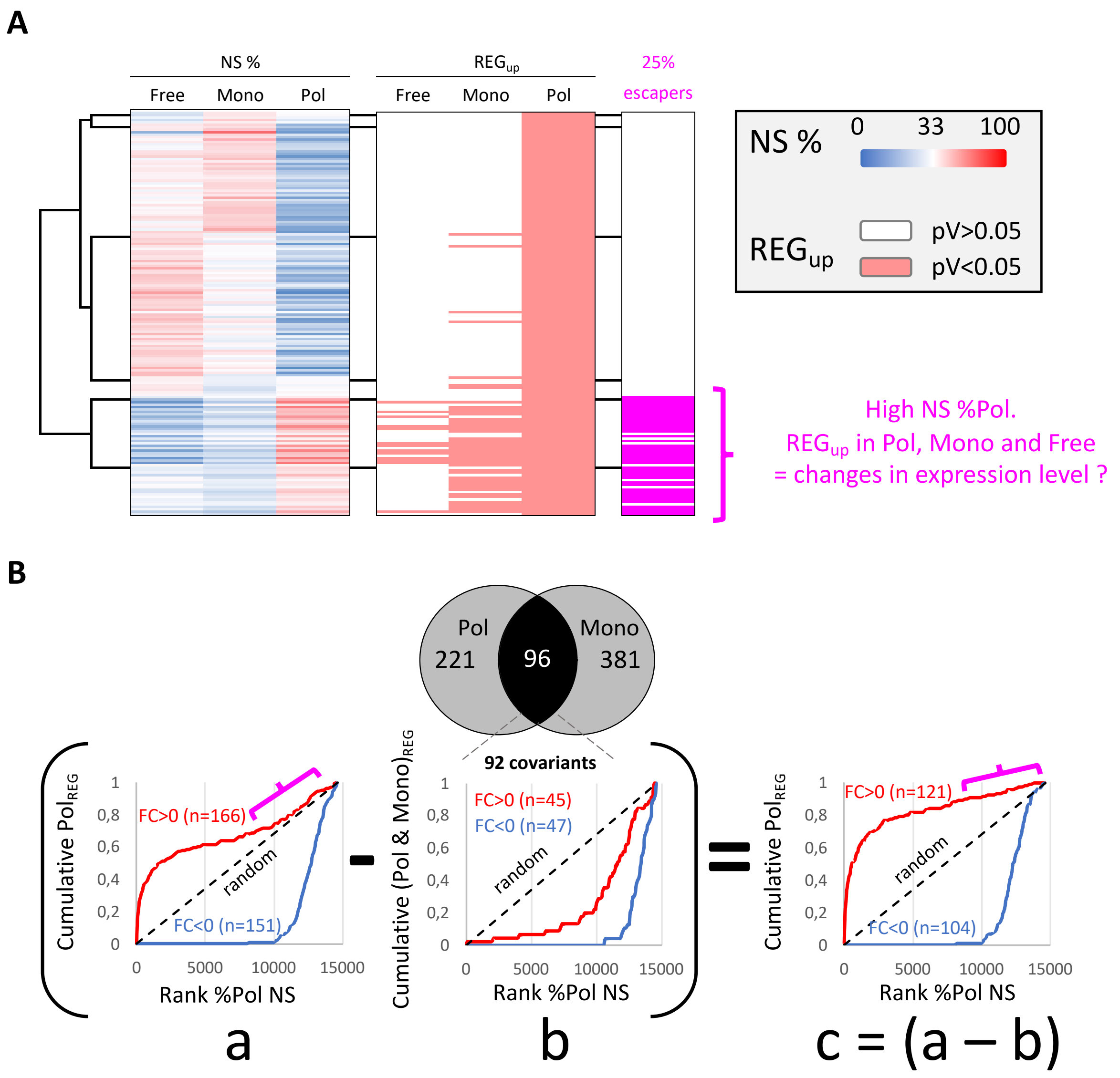


**FIGURE S5, related to Figure 5**: **Analysis of the origin of the 25% up-regulated Pol$_{REG}$ "escapers".** (**A**) Hierarchical clustering of the 166 up-regulated Pol$_{REG}$ mRNAs based on their basal distribution in Free, Mono and Pol (NS % clustering). FSH up-regulated mRNAs in each pool are indicated in the REG$_{up}$ panel. The last two clusters are mRNAs with high enrichment level in Pol compared to Mono and Free (NS % clustering, red color code in Pol and blue color code in Free and Mono). These two clusters turn out to contain most of mRNAs co-up-regulated in Pol and Mono and all mRNAs co-up-regulated in the three fractions. They also contain the top 25% mRNAs the most enriched in Pol before stimulation (25% escapers panel). (**B**) Cumulative distributions of the Pol$_{REG}$ that are not co-up or co-down regulated in Mono (right plot), based on the mRNA enrichment level in Pol before FSH stimulation (rank %Pol NS). Subtraction of the 92 positively co-up or co-down regulated mRNAs in Pol and Mono from the total 317 Pol$_{REG}$ mostly affects the right part of the cumulative distribution of the Pol$_{UP}$, corresponding mainly to the 25% escapers (pink bracket).

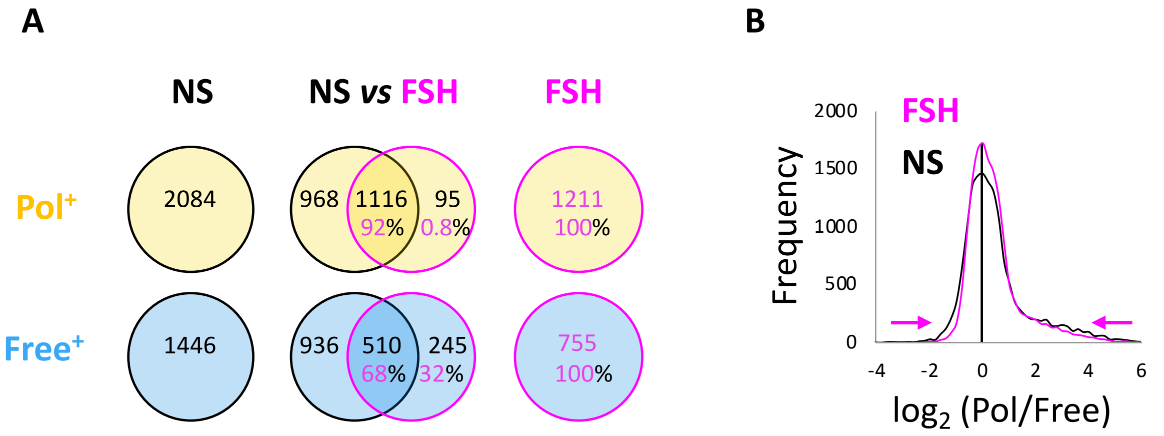


**FIGURE S6, related to Figure 6: mRNA distribution landscape remodeling by FSH.** (**A**) Comparison of mRNA content changes induced by FSH in $Pol^+$ and $Free^+$ mRNAs population. In FSH condition, 92% of $Pol^+$ and 68% of $Free^+$ mRNAs come from the basal (NS) content. (**B**) Effect of FSH on the mRNA enrichment deviations in favor of Pol or Free compared to the main distribution pattern. Pink arrows: FSH induces a shrinkage in distribution heterogeneity between mRNAs.

***Legends to supplementary tables***

**Table S1**: references of the reagents and chemicals

**Table S2**: sample description and clustering

**Table S3, related to Figure 1B**: processed data of the Sertoli cell translatome in all the conditions depicted in this study.

**Table S4, related to Figure 1C**: list of mRNAs relevant to the testicular function. The major expression site is extracted from Human Protein Atlas for single cell RNA profiles in Sertoli cells (specialized epithelial cells). 'SC' means that the mRNA is expressed only in Sertoli cells; 'SC mainly' means that the mRNA is more abundant in SC than in any other cell type; 'SC in the Testis' means that the mRNA is predominantly expressed in SC when compared to other cell types of the testis. Otherwise, the relative quantity of each mRNA in the testis is indicated. In the range column is indicated the position of Sertoli cells with respect to all other cell types that express the same mRNA, and in brackets, the number of nTPM and the percentage it represents versus the tissue where the mRNA is expressed the most (for example range 1 is 100%). Female tissues have been considered as a confounding factor, hence are not included in this analysis. In yellow are highlighted the functions of the corresponding protein when known in the testis, with the relevant references.

SC: Sertoli cell; LC: Leydig cell; Spg: spermatogonia; Spd: spermatid; Spc: spermatocyte; GC: germ cell; GPCR: G protein-coupled receptor; BTB: blood-testis barrier; TF: transcription factor; ADCC: antigen-dependent cell cytotoxicity; ECM: extra-cellular matrix; TKR: Tyrosine kinase receptor; STK: Serine/Threonine kinase; nd: not determined

**Table S5, related to Figure 2**: lists of mRNAs enriched or depleted in basal conditions in each of the pools analyzed here, namely Pol, Mono and Free.

**Table S6, related to Figure 3**: list of mRNAs down- and up-regulated in each pool, upon FSH stimulation.

**Table S7, related to Figure 5D**: lists of mRNAs that are not only highly present in Pol in resting conditions, but also exhibit increased polysomal recruitment upon FSH stimulation ("escapers")

***Supplementary references to Table S4***

An, J., Zhang, X., Qin, J., Wan, Y., Hu, Y., Liu, T., Li, J., Dong, W., Du, E., Pan, C., Zeng, W., 2014. The histone methyltransferase ESET is required for the survival of spermatogonial stem/progenitor cells in mice. Cell Death Dis. 5, e1196–e1196. https://doi.org/10.1038/cddis.2014.171

Andersen, O.M., Yeung, C.-H., Vorum, H., Wellner, M., Andreassen, T.K., Erdmann, B., Mueller, E.-C., Herz, J., Otto, A., Cooper, T.G., Willnow, T.E., 2003. Essential Role of the Apolipoprotein E Receptor-2 in Sperm Development. J. Biol. Chem. 278, 23989–23995. https://doi.org/10.1074/jbc.M302157200

Basciani, S., De Luca, G., Dolci, S., Brama, M., Arizzi, M., Mariani, S., Rosano, G., Spera, G., Gnessi, L., 2008. Platelet-derived growth factor receptor beta-subtype regulates proliferation and migration of gonocytes. Endocrinology 149, 6226–6235. https://doi.org/10.1210/en.2008-0349

Boyer, A., Lussier, J.G., Sinclair, A.H., McClive, P.J., Silversides, D.W., 2004. Pre-Sertoli Specific Gene Expression Profiling Reveals Differential Expression of Ppt1 and Brd3 Genes Within the Mouse Genital Ridge at the Time of Sex Determination1. Biol. Reprod. 71, 820–827. https://doi.org/10.1095/biolreprod.104.029371

Carré, G.-A., Couty, I., Hennequet-Antier, C., Govoroun, M.S., 2011. Gene Expression Profiling Reveals New Potential Players of Gonad Differentiation in the Chicken Embryo. PLoS ONE 6, e23959. https://doi.org/10.1371/journal.pone.0023959

Chauvin, T.R., Griswold, M.D., 2004. Characterization of the Expression and Regulation of Genes Necessary for myo-Inositol Biosynthesis and Transport in the Seminiferous Epithelium1. Biol. Reprod. 70, 744–751. https://doi.org/10.1095/biolreprod.103.022731

Chen, S.-R., Chen, M., Wang, X.-N., Zhang, J., Wen, Q., Ji, S.-Y., Zheng, Q.-S., Gao, F., Liu, Y.-X., 2013. The Wilms Tumor Gene, Wt1, Maintains Testicular Cord Integrity by Regulating the Expression of Col4a1 and Col4a21. Biol. Reprod. 88. https://doi.org/10.1095/biolreprod.112.105379

Cupp, A.S., Uzumcu, M., Skinner, M.K., 2003. Chemotactic Role of Neurotropin 3 in the Embryonic Testis That Facilitates Male Sex Determination1. Biol. Reprod. 68, 2033–2037. https://doi.org/10.1095/biolreprod.102.012617

De Santa Barbara, P., Bonneaud, N., Boizet, B., Desclozeaux, M., Moniot, B., Sudbeck, P., Scherer, G., Poulat, F., Berta, P., 1998. Direct interaction of SRY-related protein SOX9 and steroidogenic factor 1 regulates transcription of the human anti-Müllerian hormone gene. Mol. Cell. Biol. 18, 6653–6665. https://doi.org/10.1128/MCB.18.11.6653

Di-Luoffo, M., Brousseau, C., Bergeron, F., Tremblay, J.J., 2015. The Transcription Factor MEF2 Is a Novel Regulator of Gsta Gene Class in Mouse MA-10 Leydig Cells. Endocrinology 156, 4695–4706. https://doi.org/10.1210/en.2015-1500

El-Gehani, F., Tena-Sempere, M., Ruskoaho, H., Huhtaniemi, I., 2001. Natriuretic peptides stimulate steroidogenesis in the fetal rat testis. Biol. Reprod. 65, 595–600. https://doi.org/10.1095/biolreprod65.2.595

Ford, S.L., Reinhart, A.J., Lukyanenko, Y., Hutson, J.C., Stocco, D.M., 1999. Pregnenolone synthesis in immature rat Sertoli cells. Mol. Cell. Endocrinol. 157, 87–94. https://doi.org/10.1016/s0303-7207(99)00155-0

George, R.M., Hahn, K.L., Rawls, A., Viger, R.S., Wilson-Rawls, J., 2015. Notch signaling represses GATA4-induced expression of genes involved in steroid biosynthesis. REPRODUCTION 150, 383–394. https://doi.org/10.1530/REP-15-0226

Gow, A., Southwood, C.M., Li, J.S., Pariali, M., Riordan, G.P., Brodie, S.E., Danias, J., Bronstein, J.M., Kachar, B., Lazzarini, R.A., 1999. CNS myelin and sertoli cell tight junction strands are absent in Osp/claudin-11 null mice. Cell 99, 649–659. https://doi.org/10.1016/s0092-8674(00)81553-6

He, J., Li, J., Li, Y., Xu, Z., Ma, M., Chen, H., Chen, P., Lv, L., Shang, X., Liu, G., 2024. Single-cell transcriptomics identifies senescence-associated secretory phenotype (SASP) features of testicular aging in human. Aging 16, 3350–3362. https://doi.org/10.18632/aging.205538

Kanatsu-Shinohara, M., Morimoto, H., Shinohara, T., 2012. Enrichment of Mouse Spermatogonial Stem Cells by Melanoma Cell Adhesion Molecule Expression1. Biol. Reprod. 87. https://doi.org/10.1095/biolreprod.112.103861

Konrad, L., Albrecht, M., Renneberg, H., Ulrix, W., Hoeben, E., Verhoeven, G., Aumüller, G., 2000. Mesenchymal entactin-1 (nidogen-1) is required for adhesion of peritubular cells of the rat testis in vitro. Eur. J. Cell Biol. 79, 112–120. https://doi.org/10.1078/S0171-9335(04)70013-8

Kroft, T.L., Patterson, J., Won Yoon, J., Doglio, L., Walterhouse, D.O., Iannaccone, P.M., Goldberg, E., 2001. GLI1 Localization in the Germinal Epithelial Cells Alternates Between Cytoplasm and Nucleus: Upregulation in Transgenic Mice Blocks Spermatogenesis in Pachytene1. Biol. Reprod. 65, 1663–1671. https://doi.org/10.1095/biolreprod65.6.1663

Liu, C., Rodriguez, K., Yao, H.H.-C., 2016. Mapping lineage progression of somatic progenitor cells in the mouse fetal testis. Development 143, 3700–3710. https://doi.org/10.1242/dev.135756

Livera, G., Xie, F., Garcia, M.A., Jaiswal, B., Chen, J., Law, E., Storm, D.R., Conti, M., 2005. Inactivation of the mouse adenylyl cyclase 3 gene disrupts male fertility and spermatozoon function. Mol. Endocrinol. Baltim. Md 19, 1277–1290. https://doi.org/10.1210/me.2004-0318

Lu, Q., Gore, M., Zhang, Q., Camenisch, T., Boast, S., Casagranda, F., Lai, C., Skinner, M.K., Klein, R., Matsushima, G.K., Earp, H.S., Goff, S.P., Lemke, G., 1999. Tyro-3 family receptors are essential regulators of mammalian spermatogenesis. Nature 398, 723–728. https://doi.org/10.1038/19554
Matzuk, M.M., Finegold, M.J., Su, J.-G.J., Hsueh, A.J.W., Bradley, A., 1992. α-Inhibin is a tumour-suppressor gene with gonadal specificity in mice. Nature 360, 313–319. https://doi.org/10.1038/360313a0

Merkin, J., Russell, C., Chen, P., Burge, C.B., 2012. Evolutionary Dynamics of Gene and Isoform Regulation in Mammalian Tissues. Science 338, 1593–1599. https://doi.org/10.1126/science.1228186

Münsterberg, A., Lovell-Badge, R., 1991. Expression of the mouse anti-müllerian hormone gene suggests a role in both male and female sexual differentiation. Dev. Camb. Engl. 113, 613–624. https://doi.org/10.1242/dev.113.2.613

Nef, S., Verma-Kurvari, S., Merenmies, J., Vassalli, J.-D., Efstratiadis, A., Accili, D., Parada, L.F., 2003. Testis determination requires insulin receptor family function in mice. Nature 426, 291–295. https://doi.org/10.1038/nature02059

Pelletier, R.-M., Akpovi, C.D., Chen, L., Kumar, N.M., Vitale, M.L., 2015. Complementary expression and phosphorylation of Cx46 and Cx50 during development and following gene deletion in mouse and in normal and orchitic mink testes. Am. J. Physiol.-Regul. Integr. Comp. Physiol. 309, R255–R276. https://doi.org/10.1152/ajpregu.00152.2015

Perrard, M.-H., Vigier, M., Damestoy, A., Chapat, C., Silandre, D., Rudkin, B.B., Durand, P., 2007. beta-Nerve growth factor participates in an auto/paracrine pathway of regulation of the meiotic differentiation of rat spermatocytes. J. Cell. Physiol. 210, 51–62. https://doi.org/10.1002/jcp.20805

Qi, H., Deng, Z., Ye, F., Gou, J., Huang, M., Xiang, H., Li, H., 2024. Analysis of the differentially expressed genes in the combs and testes of Qingyuan partridge roosters at different developmental stages. BMC Genomics 25, 33. https://doi.org/10.1186/s12864-024-09960-2

Raymond, A.S., Shur, B.D., 2009. A novel role for SED1 (MFG-E8) in maintaining the integrity of the epididymal epithelium. J. Cell Sci. 122, 849–858. https://doi.org/10.1242/jcs.041731

Sağraç, D., Şenkal, S., Hayal, T.B., Demirci, S., Şişli, H.B., Asutay, A.B., Doğan, A., 2022. Protective role of Cytoglobin and Neuroglobin against the Lipopolysaccharide (LPS)-induced inflammation in Leydig cells ex vivo. Reprod. Biol. 22, 100595. https://doi.org/10.1016/j.repbio.2021.100595

Sarraj, M.A., Escalona, R.M., Umbers, A., Chua, H.K., Small, C., Griswold, M., Loveland, K., Findlay, J.K., Stenvers, K.L., 2010. Fetal Testis Dysgenesis and Compromised Leydig Cell Function in Tgfbr3 (Betaglycan) Knockout Mice1. Biol. Reprod. 82, 153–162. https://doi.org/10.1095/biolreprod.109.078766

Schrade, A., Kyrönlahti, A., Akinrinade, O., Pihlajoki, M., Fischer, S., Rodriguez, V.M., Otte, K., Velagapudi, V., Toppari, J., Wilson, D.B., Heikinheimo, M., 2016. GATA4 Regulates Blood-Testis Barrier Function and Lactate Metabolism in Mouse Sertoli Cells. Endocrinology 157, 2416–2431. https://doi.org/10.1210/en.2015-1927

Shen, Y., Shami, A.N., Moritz, L., Larose, H., Manske, G.L., Ma, Q., Zheng, X., Sukhwani, M., Czerwinski, M., Sultan, C., Chen, H., Gurczynski, S.J., Spence, J.R., Orwig, K.E., Tallquist, M., Li, J.Z., Hammoud, S.S., 2021. TCF21+ mesenchymal cells contribute to testis somatic cell development, homeostasis, and regeneration in mice. Nat. Commun. 12, 3876. https://doi.org/10.1038/s41467-021-24130-8

Shiromoto, Y., Kuramochi-Miyagawa, S., Nagamori, I., Chuma, S., Arakawa, T., Nishimura, T., Hasuwa, H., Tachibana, T., Ikawa, M., Nakano, T., 2019. GPAT2 is required for piRNA biogenesis, transposon silencing, and maintenance of spermatogonia in mice†. Biol. Reprod. 101, 248–256. https://doi.org/10.1093/biolre/ioz056

Silva, E.J.R., Patrão, M.T.C.C., Tsuruta, J.K., O'Rand, M.G., Avellar, M.C.W., 2012. Epididymal protease inhibitor (EPPIN) is differentially expressed in the male rat reproductive tract and immunolocalized in maturing spermatozoa. Mol. Reprod. Dev. 79, 832–842. https://doi.org/10.1002/mrd.22119

Sun, X., Wang, Y., Mai, Q., Fan, Y., Huang, Q., Du, Y., Huang, H., Ma, E., Wang, F., Zhang, G., 2023. Neuromedin S regulates steroidogenesis and proliferation in goat Leydig cells through modulating mitochondrial function. FASEB J. 37, e22989. https://doi.org/10.1096/fj.202300187R

Szczepny, A., Hogarth, C.A., Young, J., Loveland, K.L., 2009. Identification of Hedgehog Signaling Outcomes in Mouse Testis Development Using a Hanging Drop-Culture System1. Biol. Reprod. 80, 258–263. https://doi.org/10.1095/biolreprod.108.067926

Tsuchiya, H., Fujinoki, M., Azuma, M., Koshimizu, T., 2023. Vasopressin V1a receptor and oxytocin receptor regulate murine sperm motility differently. Life Sci. Alliance 6, e202201488. https://doi.org/10.26508/lsa.202201488

Uchida, A., Sakib, S., Labit, E., Abbasi, S., Scott, W., Underhill, M., Biernaskie, J., Dobrinski, I., 2020. Development and function of smooth muscle cells is modulated by *Hic1* in mouse testis. Development dev.185884. https://doi.org/10.1242/dev.185884

Uhrin, P., Dewerchin, M., Hilpert, M., Chrenek, P., Schöfer, C., Zechmeister-Machhart, M., Krönke, G., Vales, A., Carmeliet, P., Binder, B.R., Geiger, M., 2000. Disruption of the protein C inhibitor gene results in impaired spermatogenesis and male infertility. J. Clin. Invest. 106, 1531–1539. https://doi.org/10.1172/JCI10768

Wang, M.-W., Yang, Z., Chen, X., Zhou, S.-H., Huang, G.-L., Sun, J.-N., Jiang, H., Xu, W.-M., Lin, H.-C., Yu, X., Sun, J.-P., 2021. Activation of PTH1R alleviates epididymitis and orchitis through Gq and β-arrestin-1 pathways. Proc. Natl. Acad. Sci. 118, e2107363118. https://doi.org/10.1073/pnas.2107363118

Wang, Y., Li, T., Qiu, X., Mo, X., Zhang, Y., Song, Q., Ma, D., Han, W., 2008. CMTM3 can affect the transcription activity of androgen receptor and inhibit the expression level of PSA in LNCaP cells. Biochem. Biophys. Res. Commun. 371, 54–58. https://doi.org/10.1016/j.bbrc.2008.03.143

Xu, J., Sang, M., Cheng, J., Luo, C., Shi, J., Sun, F., 2023. Knockdown of disheveled-associated activator of morphogenesis 2 disrupts cytoskeletal organization and phagocytosis in rat Sertoli cells. Mol. Cell. Endocrinol. 563, 111867. https://doi.org/10.1016/j.mce.2023.111867

Yao, S., Wei, X., Deng, W., Wang, B., Cai, J., Huang, Y., Lai, X., Qiu, Y., Wang, Y., Guan, Y., Wang, J., 2022. Nestin-dependent mitochondria-ER contacts define stem Leydig cell differentiation to attenuate male reproductive ageing. Nat. Commun. 13, 4020. https://doi.org/10.1038/s41467-022-31755-w

Yin, Y., Wang, G., Liang, N., Zhang, H., Liu, Z., Li, W., Sun, F., 2013. Nuclear export factor 3 is involved in regulating the expression of TGF-β3 in an mRNA export activity-independent manner in mouse Sertoli cells. Biochem. J. 452, 67–78. https://doi.org/10.1042/BJ20121006

Zeng, F., Zhu, X., Li, C., Han, B., Meng, L., Li, L., Wei, H., Zhang, S., 2022. Carboxypeptidase E protein regulates porcine sperm Ca2+ influx to affect capacitation and fertilization. Theriogenology 192, 28–37. https://doi.org/10.1016/j.theriogenology.2022.08.017
